# The spatial extent of anatomical connections within the thalamus varies across the cortical hierarchy in humans and macaques

**DOI:** 10.1101/2023.07.22.550168

**Authors:** Amber M. Howell, Shaun Warrington, Clara Fonteneau, Youngsun T. Cho, Stamatios N. Sotiropoulos, John D. Murray, Alan Anticevic

## Abstract

Each cortical area has a distinct pattern of anatomical connections within the thalamus, a central subcortical structure composed of functionally and structurally distinct nuclei. Previous studies have suggested that certain cortical areas may have more extensive anatomical connections that target multiple thalamic nuclei, which potentially allows them to modulate distributed information flow. However, there is a lack of quantitative investigations into anatomical connectivity patterns within the thalamus. Consequently, it remains unknown if cortical areas exhibit systematic differences in the extent of their anatomical connections within the thalamus. To address this knowledge gap, we used diffusion magnetic resonance imaging (dMRI) to perform brain-wide probabilistic tractography for 828 healthy adults from the Human Connectome Project. We then developed a framework to quantify the spatial extent of each cortical area’s anatomical connections within the thalamus. Additionally, we leveraged resting-state functional MRI, cortical myelin, and human neural gene expression data to test if the extent of anatomical connections within the thalamus varied along the cortical hierarchy, from sensory and motor to multimodal associative cortical areas. Our results revealed two distinct corticothalamic tractography motifs: 1) a sensorimotor cortical motif characterized by focal thalamic connections targeting posterolateral thalamus, associated with fast, feed-forward information flow; and 2) an associative cortical motif characterized by diffuse thalamic connections targeting anteromedial thalamus, associated with slow, feed-back information flow. These findings were consistent across human subjects and were also observed in macaques, indicating cross-species generalizability. Overall, our study demonstrates that sensorimotor and association cortical areas exhibit differences in the spatial extent of their anatomical connections within the thalamus, which may support functionally-distinct cortico-thalamic information flow.

## Introduction

Mapping the anatomical connections of the brain is a fundamental goal in neuroscience, as these pathways tether brain areas together and impose constraints on their functional interactions. The thalamus is a central subcortical structure that is extensively connected to the entire cortex through long-range white matter fiber tracts (1). These tracts form parallel cortico-thalamic circuits, which enable the thalamus to relay and coordinate information across the cortex (2–4). Notably, the thalamus lacks reciprocal excitatory connections (5). Thus, computations involving the thalamus rely on its long-range inputs from and outputs to the cortex (6). Mapping these long-range connections can provide insight into the role of the thalamus in shaping cortical information flow and the neural basis of cognitive computation, both of which are critically reliant on the interactions between the thalamus and cortex in vertebrates (2, 7–12).

Studies of cortical-thalamic connectivity date back to the early 19th century, yet we still lack a comprehensive understanding of how these connections are organized (see 13 and 14 for review). The traditional view of the thalamus is based on its histologically-defined nuclear structure (6). This view was originally supported by evidence that cortical areas project to individual thalamic nuclei, suggesting that the thalamus primarily relays information (15). However, several studies have demonstrated that cortical connectivity within the thalamus is topographically organized and follows a smooth gradient across the thalamus (16–21). Additionally, some cortical areas exhibit extensive connections within the thalamus, which target multiple thalamic nuclei (22, 23). These extensive connections may enable information integration within the thalamus through overlapping termination patterns from different cortical areas, a key mechanism for higher-order associative thalamic computations (24–26). However, our knowledge of how thalamic connectivity patterns vary across cortical areas, especially in humans, remains incomplete. Characterizing cortical variation in thalamic connectivity patterns may offer insights into the functional roles of distinct cortico-thalamic loops (6, 7).

Primate studies investigating cortico-thalamic circuitry have primarily relied on anatomical tracer data in monkeys (e.g., 18–20, 27–29). However, such invasive studies cannot be replicated in humans. Fortunately, advancements in magnetic resonance imaging (MRI) have enabled the examination of white matter tracts *in vivo* using diffusion MRI (dMRI), which measures the diffusion properties of water molecules in brain tissue (30–32). These properties are then used by tractography algorithms to reconstruct white matter tracts, known as streamlining, and estimate region-to-region connectivity (33–36).

State-of-the-art tractography techniques can now map streamlines at high spatial resolutions to reveal connectivity patterns within the thalamus (16, 22, 23, 37, 38). These studies have unveiled a diverse array of cortico-thalamic circuits, which have unique origins, targets, strengths, and microstructural profiles and are optimized for distinct roles in sensory and higher-order associative computations (22, 23, 39–41). Emerging dMRI evidence also suggests that certain cortical areas may have more extensive connections within the thalamus (23), which may grant them privileged access for integrating cortical signals or modulating whole-brain functional interactions (42). However, quantitative DWI studies that examine the spatial patterns of brain connections are limited (36). Therefore, it is uncertain whether the extent of anatomical connections within the thalamus systematically varies across cortical areas in humans and what implications such variation may have for information processing within distinct cortico-thalamic systems.

Furthermore, there are few studies that have directly compared cortico-thalamic anatomical circuitry between humans and non-human primates using tractography methods (e.g., 41, 43). Such studies form a bridge to the existing macaque tract tracing literature. They also provide validation for human dMRI findings, as macaque dMRI can be collected at much higher resolutions, without confounds such as motion artifacts (44).

While previous work has demonstrated that some thalamic territories diffusely project across the cortex, little work has been done to examine spatial patterns of connections within the thalamus. The aim of this study was to investigate the spatial extent of anatomical connectivity patterns within the thalamus in both humans and non-human primates and determine if such patterns differ between sensorimotor and association cortical areas. To this end, we leveraged probabilistic tractography derived from 3T diffusion data from 828 healthy human adults from the Human Connectome Project (HCP) and 7T diffusion data from six post-mortem macaque monkeys. We first developed an approach to quantify the spatial properties of anatomical connectivity patterns within the thalamus. We then tested if the extent of these patterns varied across the cortical hierarchy, to determine if the extent of these patterns was different between sensorimotor and associative cortical areas. We found that sensorimotor cortical areas exhibited more focal thalamic connectivity patterns, while association cortical areas exhibited more diffuse, or extensive, thalamic connectivity patterns. Additionally, we show that such cortical variation was consistent across individuals, generalized in macaques, and associated with distinct types of information flow. Overall, our findings highlight that sensorimotor and association cortical areas exhibit distinct anatomical connectivity patterns within the thalamus, and differences in the extent of such thalamic connectivity patterns may support functionally distinct cortico-thalamic computations.

## Results

### Cortical areas differ in the extent of their anatomical connectivity patterns within the thalamus

To test if there are systematic differences in the spatial extent of anatomical connections within the thalamus across cortical areas, we developed a framework to quantify the spatial extent of thalamic connectivity patterns using Euclidean distance (ED) **(Fig. 1A)**. This framework assigns a value to every cortical area (referred to throughout the manuscript as *ED*_*pc*1_ loadings). This measure reflects the spatial extent of each cortical area’s anatomical connections within the thalamus, such that cortical areas with higher *ED*_*pc*1_ loadings have more focal thalamic connections.

**Fig. 1.**
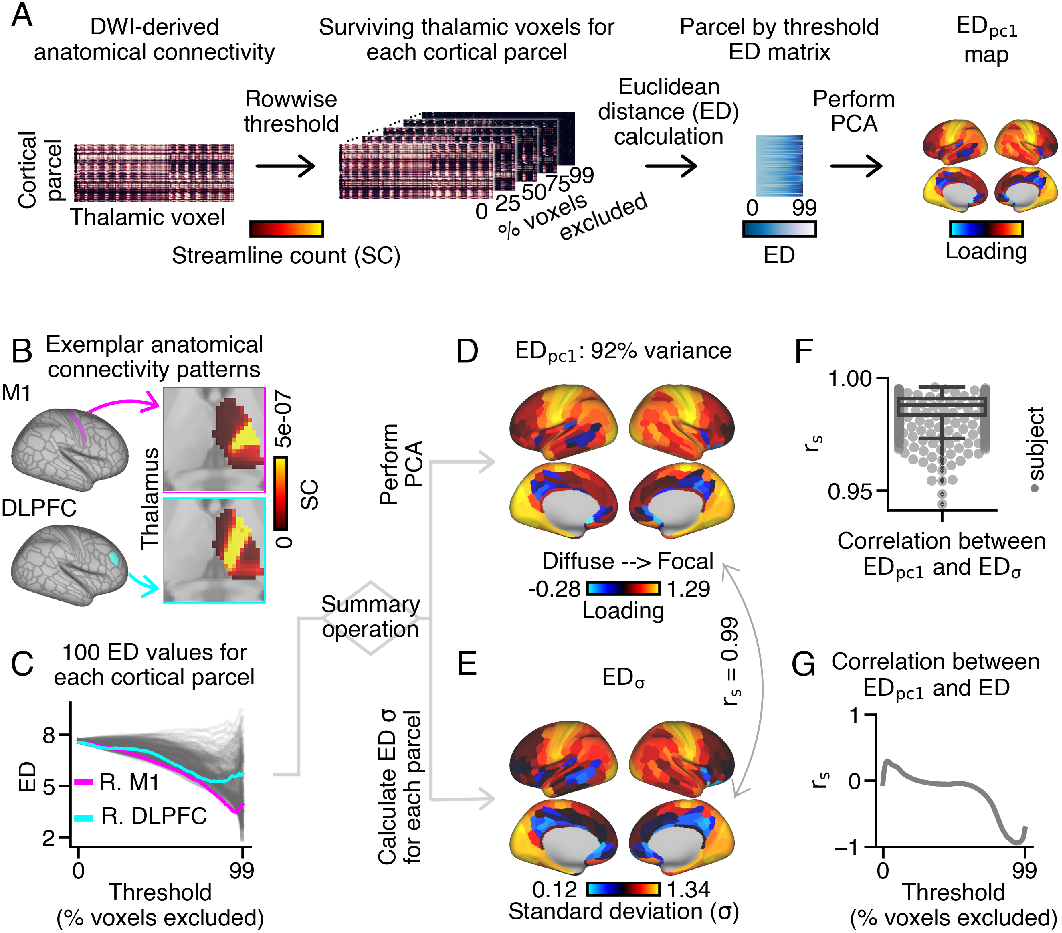
Workflow to quantify the extent of each cortical area’s thalamic anatomical connectivity pattern using Euclidean distance (ED). **(A)** Schematic overview of the thresholding and ED calculation framework applied to group-averaged and individual-level human probabilistic tractography data (n=828). ED was used to measure the extent of each cortical area’s anatomical connectivity pattern within ipsilateral thalamus (see **Fig. S7** for bilateral calculation). **(B)** Thalamic connectivity patterns for right motor area 1 (M1) and dorsolateral prefrontal cortex (DLPFC). M1 and DLPFC exhibit focal and diffuse thalamic connections, respectively. Cortical parcels were defined via Glasser et al. (45). **(C)** ED matrix, depicting average pairwise distances between surviving thalamic voxels for 100 thresholds. Cortical parcels with more diffuse thalamic connections exhibited higher ED values across thresholds (e.g., DLPFC). **(D)** Cortical *ED*_*pc*1_ loading map obtained by applying principal component analysis (PCA) to the ED matrix. **(E)** Cortical *ED*_*σ*_ map showing the standard deviation of ED across 100 thresholds for each cortical parcel. *ED*_*σ*_ almost perfectly correlated with *ED*_*pc*1_ (*r*_*s*_=0.99; *p*_*sa*_ < 0.001), with p-values estimated using spatial-autocorrelation (sa) preserving surrogate brain maps (46). *r*_*s*_ : Spearman rho. **(F)** Correlations between *ED*_*pc*1_ and *ED*_*σ*_ were highly consistent across subjects. **(G)** *ED*_*pc*1_ loadings and ED values across cortex negatively correlated at more conservative thresholds, indicating that higher *ED*_*pc*1_ loadings correspond to more focal thalamic connections.

Briefly, the framework uses anatomical connectivity between each cortical area and each thalamic voxel, creating a matrix where each element is a streamline count (SC). A higher streamline count reflects a higher likelihood that a thalamic voxel is connected to a given cortical area relative to lower streamline counts. Because we used probabilistic tractography, which assigns a probabilistic estimate to each thalamic voxel, each cortical area’s thalamic streamline counts must be thresholded to exclude thalamic voxels with low streamline counts. There is no consensus for selecting a threshold, so each cortical area’s thalamic streamline counts were iteratively thresholded by excluding 0% to 99% of thalamic voxels with the lowest streamline counts. Next, average pairwise ED was calculated between these ‘surviving’ thalamic voxels (i.e., the top x% of thalamic voxels with the highest streamline counts) for each threshold. This resulted in a matrix of 100 ED values for each cortical area. This ‘ED matrix’ was then used as input into a principal components analysis (PCA) to produce a single loading value for each cortical area (*ED*_*pc*1_ loadings). We show that *ED*_*pc*1_ loadings index the spatial extent of each cortical area’s anatomical connections within the thalamus, such that a cortical areas with higher *ED*_*pc*1_ loadings have more focal thalamic connections.

We applied the thresholding and ED framework to whole-brain dMRI-derived probabilistic tractography data from healthy adults from the Human Connectome Project (n=828; see SI Appendix **Fig. S1** for preprocessing steps). We segmented the cortex into 360 discrete areas using cortical parcels defined in the Glasser et al. parcellation (47) and extracted SCs between these cortical parcels and each thalamic voxel. Only ipsilateral thalamic voxels were considered, unless otherwise specified in supplementary analyses. Each cortical parcel exhibited a distinct thalamic connectivity pattern, as exemplified by motor area 1 (M1; top magenta panel) and dorsolateral prefrontal cortex (DLPFC; bottom cyan panel) **(Fig. 1B)**. Here, warmer colors reflect voxels that have a higher likelihood of an anatomical connection to the given cortical area relative to cooler colors.

Next, for each cortical parcel, we iteratively calculated ED between thalamic voxels after progressively excluding thalamic voxels with the lowest SCs (**Fig. S2** shows the surviving thalamic voxels for a subset of thresholds). This process generated 100 ED values for 360 cortical parcels and 100 thresholds **(Fig. 1C)**. As thalamic voxels with lower SCs were progressively excluded from the ED calculation, cortical areas with more diffuse thalamic connections had ED values that remain relatively higher across thresholds (e.g., DLPFC).

We then conducted PCA using the ED matrix as input to derive a single value for each cortical parcel. The first PC accounted for almost all of the variation of ED across cortical parcels (92%) **(Fig. 1D)**. Cortical parcels with the lowest loadings on PC1 (*ED*_*pc*1_ loadings) included bilateral anterior cingulate areas, bilateral precuneus, and right temporal cortex. In contrast, cortical parcels with the highest *ED*_*pc*1_ loadings included bilateral visual areas, right somatosensory cortex, and left entorhinal cortex (refer to **Fig. S3** for visualizations of their respective thalamic connectivity patterns).

We then compared *ED*_*pc*1_ loadings with measures derived from alternative methods to quantify the extent of cortical connections within the thalamus. Based on the qualitative observation that more conservative thresholds showed the most variation in ED values across cortical parcels, the standard deviation of ED across thresholds was calculated for each cortical parcel (*ED*_*σ*_) **(Fig. 1E)**. Remarkably, the *ED*_*pc*1_ and *ED*_*σ*_ measures almost perfectly correlated (*r*_*s*_=0.99; *p*_*sa*_ *<* 0.001), with statistical significance determined using spatial-auto-correlation (SA) preserving surrogate maps (see SI Appendix for further details) (46). The high correlation between *ED*_*pc*1_ and *ED*_*σ*_ was consistently observed across subjects (mean=0.99, max=0.99, min=0.94) **(Fig. 1F)**. This suggests the the dominant PC axis reflects ED variation at more conservative thresholds. Moreover, we explored additional measures to capture the extent of cortical connections within the thalamus, which largely replicated the main findings of this study (**Fig. S4**). Based on its superior agreement with ED calculated at more conservative thresholds, we selected *ED*_*pc*1_ loadings for further analysis.

We next examined whether the *ED*_*pc*1_ measure could distinguish focal and diffuse thalamic connections. To do this, we compared *ED*_*pc*1_ loadings with ED values calculated at individual thresholds **(Fig. 1G)**. We observed a strong negative correlation between *ED*_*pc*1_ and ED at more conservative thresholds (e.g., 78-99%; **Fig. S5**). Additionally, we investigated the relationship between *ED*_*pc*1_ loadings and the mean and standard deviation of ED across threshold ranges. These measures also highly correlated with *ED*_*pc*1_ loadings at more conservative thresholds **(Fig. S6)**. Overall, higher *ED*_*pc*1_ loadings correspond to lower ED values, when calculated between thalamic voxels with the highest SCs, which reflect cortical parcels with more focal thalamic connectivity patterns.

We also performed supplementary analysis to determine if the extent of cortical connections within the thalamus varied across hemispheres by calculating *ED*_*pc*1_ loadings for bilateral **(Fig. S7)** and contralateral **(Fig. S8)** thalamic connectivity patterns. Bilateral *ED*_*pc*1_ loadings strongly correlated with ipsilateral loadings (*r*_*s*_=0.91), while contralateral loadings exhibited a weaker, yet still significant, correlation with ipsilateral loadings (*r*_*s*_=0.57). Furthermore, bilateral *ED*_*pc*1_ loadings differentiated cortical parcels with unilateral and bilateral thalamic connectivity patterns **(Fig. S7F)**, demonstrating that the *ED*_*pc*1_ measure could distinguish between the extent of thalamic connections within and across hemispheres.

Finally, we conducted extensive control analyses. We found that *ED*_*pc*1_ loadings were not associated with inter-subject variation in motion **(Fig. S9)** or volumes of gray or white matter in the cortex or subcortex **(Fig. S10)**. Additionally, inter-cortical variation of *ED*_*pc*1_ loadings did not correspond with average streamline count (Mean SC) or streamline length **(Fig. S11)**, anatomical overlap with thalamic fractional anisotropy and mean diffusivity values **(Fig. S12)**, or cortical geometry, distortion, bias, and surface area **(Fig. S13)**. *ED*_*pc*1_ loadings also remained largely unchanged after accounting for size differences between the left and right thalamus **(Fig. S14)**, inter-cortical variation of cortical curvature **(Fig. S15)**, and inter-voxel variation in total connectivity strength across ipsilateral cortex **(Fig. S31)**.

Lastly, we also calculated isotropy to capture how evenly a connectivity pattern extends within the thalamus **(Fig. S16)**, which also did not correspond highly with *ED*_*pc*1_ loadings.

### Sensory cortical parcels have more focal connections within the thalamus relative to association cortical parcels

Animal studies have shown that anatomical features in cortico-thalamic circuits are hierarchically-organized, or varying between sensorimotor and association cortical areas (49–52). Therefore, we hypothesized that the spatial extent of anatomical connections within the thalamus would vary along the cortical hierarchy in humans. To test this hypothesis, we compared *ED*_*pc*1_ loadings between sensory and association cortical parcels using both network and gradient approaches **(Fig. 2)**. Overall, we found that sensory cortical parcels had more focal thalamic connections relative to association cortical parcels.

**Fig. 2.**
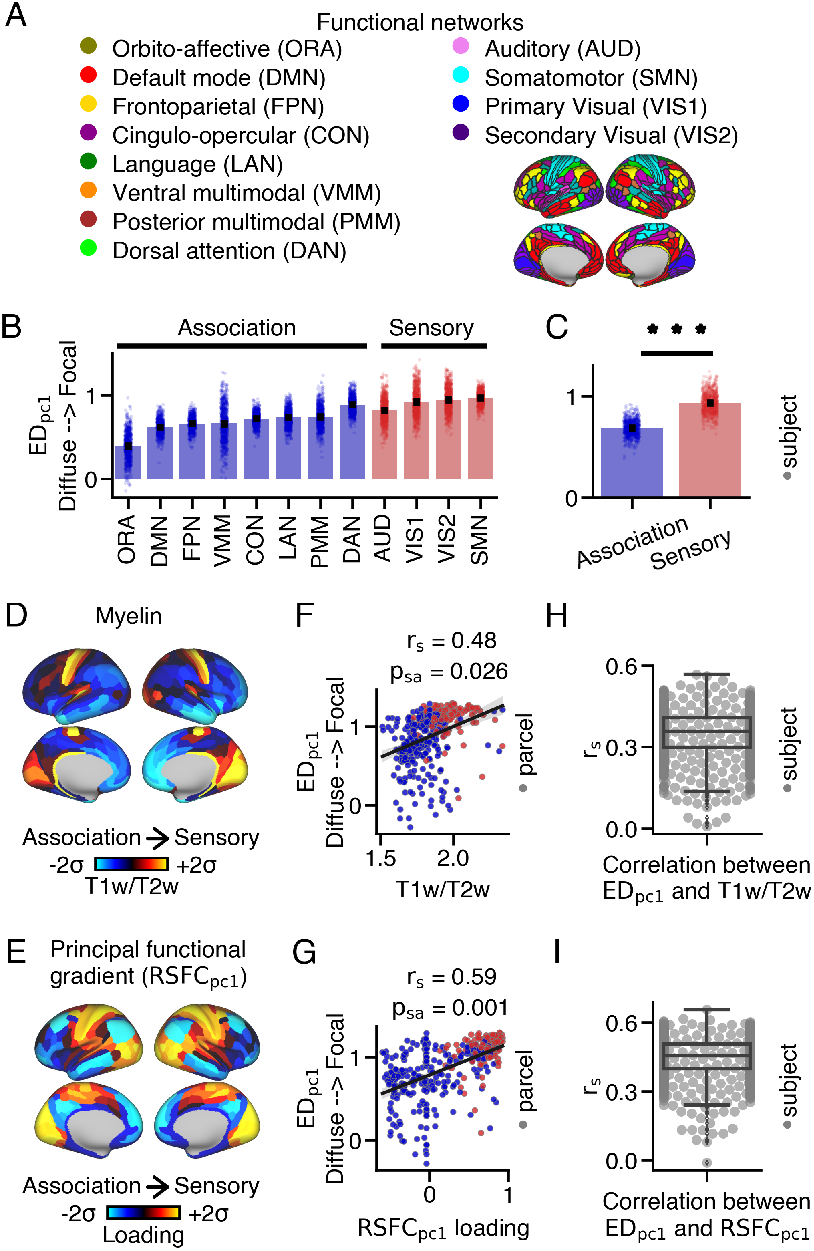
Differences in the extent of anatomical connections within the thalamus between sensory and association cortical parcels. **(A)** Resting-state functional connectivity networks identified by Ji et al. (48). **(B)** Average *ED*_*pc*1_ loading within each network for each subject. Barplots show the mean and standard error. **(C)** Sensory networks exhibited significantly higher *ED*_*pc*1_ loadings compared to association networks (two-sided Wilcoxon signed-rank test; *** p < 0.001). **(D)** Cortical myelin map calculated by averaging T1w/T2w values across subjects. **(E)** Cortical principal functional gradient (*RSFC*_*pc*1_) loading map derived from PCA on cortico-cortical resting-state functional connectivity data. Sensory cortical parcels exhibit higher T1w/T2w values and *RSFC*_*pc*1_ loadings compared to association cortical parcels. **(F**,**G)** Correlations between *ED*_*pc*1_ loadings and T1w/T2w values, as well as *RSFC*_*pc*1_ loadings, across the cortex. **(H**,**I)** On average, a moderate relationship was observed between *ED*_*pc*1_ loadings and T1w/T2w values, as well as *RSFC*_*pc*1_ loadings, across subjects. Boxplots show the median and inter-quartile ranges.

First, we assigned each cortical parcel to one of twelve resting-state functional networks based on the work of Ji et al. (48). These networks included four sensorimotor (‘sensory’) and eight higher-order associative (‘association’) networks (**Fig. 2A**). *ED*_*pc*1_ loadings were higher in sensory networks (median=0.93) compared to association networks (median=0.68) (Wilcoxon signed-rank test: W=0, p=3.76e-137) **(Fig. 2B,C)**.

Recent methods have emerged to characterize brain organization along neural gradients, which reflect smooth spatial transitions in brain features (53). Many of these gradients vary between sensory and association cortical areas, including the T1w/T2w ratio, a proxy measure of cortical myelin (54), and the principal resting-state functional gradient (*RSFC*_*pc*1_), which is derived from cortico-cortical resting-state blood-oxygen-level-dependent (BOLD) functional connectivity (55). T1w/T2w values and *RSFC*_*pc*1_ loadings highly correspond with one another (*r*_*s*_=0.51; **Fig. S17**A) and both are higher in sensory relative to association cortical parcels **(Fig. 2D,E)**.

*ED*_*pc*1_ loadings significantly correlated with both T1w/T2w values (*r*_*s*_=0.48, *p*_*sa*_=0.026) **(Fig. 2F)** and *RSFC*_*pc*1_ loadings (*r*_*s*_=0.59, *p*_*sa*_=0.001) across cortex **(Fig. 2G)**. On average, we observed a moderate correlation between each subject’s *ED*_*pc*1_ loadings and group-averaged T1w/T2w values (mean=0.35, median=0.36, SEM=0.003, STD=0.09) and group-averaged *RSFC*_*pc*1_ loading (mean=0.45, median=0.45, SEM=0.003, STD=0.09) **(Fig. 2H,I)**. However, there were instances where a weak relationship was observed between these cortical maps. These weak correlations may be attributed to the presence of biologically implausible connections originating from certain cortical parcels, which were associated with a weaker correlation between *ED*_*pc*1_ loadings and T1w/T2w values, as well as *RSFC*_*pc*1_ loadings, across subjects **(Fig. S18)**. See SI Appendix for further details.

The finding that *ED*_*pc*1_ loadings significantly correlated with T1w/T2w values, as well as *RSFC*_*pc*1_ loadings, was replicated using bilateral thalamic connectivity patterns **(Fig. S7)**, but not contralateral thalamic patterns **(Fig. S8)**. This finding was also replicated using an alternate tractography seeding strategy **(Fig. S19)**, dense connectivity data **(Fig. S20)**, and structurally and functionally defined thalamic masks **(Fig. S21)**. Lastly, at more conservative thresholds, ED calculated at individual thresholds **(Fig. S5)** and the mean and standard deviation of ED calculated for a range of thresholds **(Fig. S6)** also differed between sensory and association cortical parcels.

We also tested the correspondence between *ED*_*pc*1_ loadings and the anteroposterior cortical gradient and the secondary principal functional gradient, which reflects functional specialization along the sensory-association-motor cortical axis (55). Neither cortical gradient significantly correlated with *ED*_*pc*1_ loadings (**Fig. S22**), suggesting specificity for the sensorimotor-association cortical gradient.

### Cortical parcels with focal connections within the thalamus preferentially couple with posterolateral thalamus

Thalamic nuclei are known to play roles in sensorimotor and associative cognitive processes (9, 56). We hypothesized that cortical parcels with diffuse thalamic connections would anatomically couple with associative thalamic nuclei, while those with focal thalamic connections would anatomically couple with sensorimotor thalamic nuclei. To test this hypothesis, we compared each cortical parcel’s *ED*_*pc*1_ loadings with their anatomical coupling with thalamic nuclei associated with sensorimotor and associative computations **(Fig. 3)**. We observed that sensorimotor thalamic nuclei (e.g., posterolateral, first-order thalamic nuclei) preferentially couple with cortical parcels with focal thalamic connections. One the other hand, associative thalamic nuclei (e.g., higher-order, mediolateral thalamus) exhibit more variable targets, but overall appear to couple with cortical parcels with focal and diffuse thalamic connections.

**Fig. 3.**
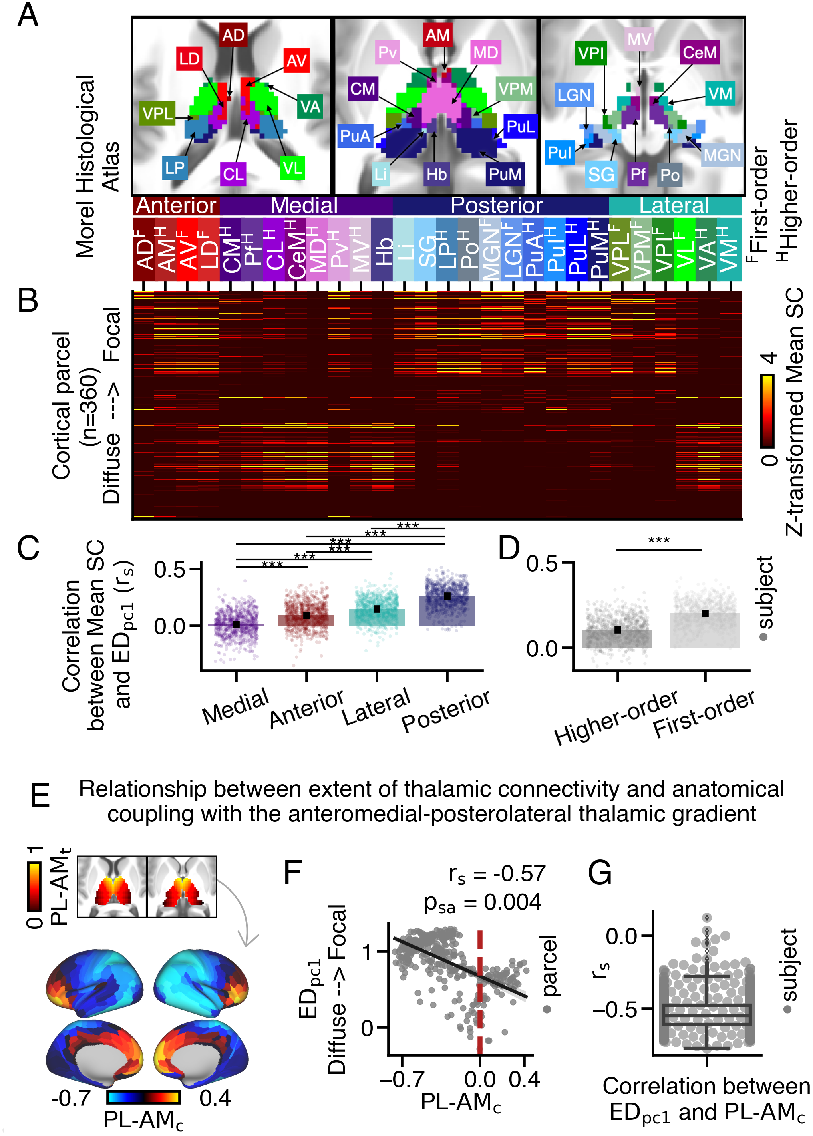
Mapping between the spatial extent of anatomical connections within the thalamus and anatomical coupling with the anteromedial-posterolateral thalamic gradient. **(A)** Axial view of 28 histologically-defined thalamic nuclei from the Morel thalamic atlas (57). **(B)** Anatomical connectivity matrix depicting the anatomical connectivity strength between each cortical parcel and each thalamic nucleus. The y-axis is sorted such that cortical parcels with higher *ED*_*pc*1_ loadings are positioned near the top. **(C-D)** The correlation between *ED*_*pc*1_ loadings and Mean SC values was calculated for each thalamic nucleus for each subject. Higher values indicate thalamic nuclei that preferentially couple with cortical parcels with focal thalamic connections. **(C)** Posterior thalamic nuclei preferentially coupled with cortical parcels with focal thalamic connections, relative to lateral, anterior, and medial nuclei (Friedman test; *** p=0.001 for Nemenyi post-hoc tests). Each dot represents a subject’s averaged *r*_*s*_ value across nuclei within the subclass. **(D)** First-order thalamic nuclei preferentially coupled with cortical parcels with focal thalamic connections (two-sided Wilcoxon signed-rank test). **(E)** The overlap between each cortical parcel’s anatomical connectivity along the anteromedial-posterolateral thalamic spatial gradient (*P L*-*AM*_*t*_). Positive values denote preferential coupling with anteromedial thalamus, while negative values denote preferential coupling with posterolateral thalamus (*P L*-*AM*_*c*_). **(F)** Cortical parcels with higher *ED*_*pc*1_ loadings preferentially coupled with the posterolateral thalamus, reflected by lower *P L*-*AM*_*c*_ values. (**G**) On average, a moderate relationship was observed between *ED*_*pc*1_ loadings and *P L*-*AM*_*c*_ values across subjects.

First, we tested if there was a correspondence between the extent and location of a cortical parcel’s connections within the thalamus. For this analysis, we used the Morel histological thalamic atlas to segment the thalamus into 28 thalamic nuclei **(Fig. 3A)**. We then constructed an anatomical connectivity matrix to assess the strength of connectivity between each cortical parcel and each thalamic nucleus **(Fig. 3B)**. In this matrix, warmer colors reflect a higher Mean SC, reflecting more likely corticothalamic connections relative to cooler colors. The cortical parcels with the highest group-averaged *ED*_*pc*1_ loadings are positioned at the top. Mean SC values were normalized within each nucleus, to better visualize patterns of anatomical connectivity across cortex parcels for each thalamic nucleus (see **Fig. S23** for unnormalized matrices).

Based on previous reports proposing different classification schemes for thalamic nuclei, we classified thalamic nuclei into subgroups based on their spatial proximity (e.g., anterior, medial, posterior, and lateral groups), primary input sources (e.g., higher/first-order; (58)), and molecular architecture (e.g., primary, secondary, and tertiary; 59; **Fig. S24)**. See **Table. S1** for all nuclei labels and subgroup assignments. Anterior and medial thalamic nuclei are commonly associated with higher-order cognitive functions, while lateral and posterior nuclei are primarily associated with sensorimotor cognitive functions (8, 42, 60, 61). Similarly, ‘first-order’ thalamic nuclei relay sensory information to the cortex. On the other hand, ‘higher-order’ thalamic nuclei facilitate trans-thalamic information processing which is important for higher-order cognitive functions (15, 58). These simplified classification schemes do not fully encompass thalamic nuclei heterogeneity, so we tested multiple classification schemes to test our hypothesis.

We correlated each cortical parcel’s *ED*_*pc*1_ loadings with Mean SC values for each thalamic nucleus, separately. This produced 28 Spearman rho values for each subject. Stronger rho values indicated thalamic nuclei that preferentially couple with cortical parcels with focal (closer to 1) or diffuse (closer to -1) connections within the thalamus. Weak rho values (closer to 0) indicated thalamic nuclei with little a weak preference between these cortical parcels (i.e., they coupling with cortical areas with focal and diffuse thalamic connectivity patterns).

We then tested if these rho values differed between thalamic nuclei. First, we found that the correlation between *ED*_*pc*1_ loadings and Mean SC was highest in posterior (median=0.27) thalamic nuclei, followed by lateral (median=0.15), anterior (median=0.1), and medial (median=0.01) thalamic nuclei (χ ^2^ (3)=1165, p < .001) **(Fig. 3C)**. Post-hoc Nemenyi tests indicated significant differences between all group comparisons (*** p=0.001). We also found that these correlations were higher in first-order (median=0.20) compared to higher-order (median=0.11) thalamic nuclei (Wilcoxon signed-rank test; W=2067, p=6.47e-134).

We also examined the correlation between *ED*_*pc*1_ loadings and Mean SC within thalamic subclasses defined based on gene expression data (59), which replicated our main findings **(Fig. S24)**. Additionally, we examined this correlation for each of the 28 Morel thalamic nuclei using group-averaged data. However, we observed modest correlations that did not survive correction for multiple comparisons **(Fig. S23)**. It is worth noting that other classifications of thalamic nuclei exist (e.g., 62), which we did not consider in this study.

Emerging studies suggest that some properties of thalamic anatomy vary along continuous spatial gradients within the thalamus (16, 21, 63). Based on previous findings that thalamic connectivity varies along anteroposterior and mediolateral spatial gradients (16, 21), we hypothesized that the extent of connections within the thalamus would also continuously vary along these axes. To investigate this hypothesis, we examined cortico-thalamic anatomical coupling along Cartesian spatial gradients within the thalamus (e.g., antero-posterior, mediolateral, dorsoventral).

First, we calculated the position of each thalamic voxel along the anteromedial-posterolateral spatial gradient (*P L*-*AM*_*t*_). Then, we correlated each cortical parcel’s streamline counts within ipsilateral thalamus to each thalamic voxel’s position along the anteromedial-posterolateral gradient (schematized in **Fig. S25**). This workflow generated a cortical map where warmer colors reflect cortical parcels that anatomically couple with anteromedial thalamus and cooler colors reflect cortical parcels that anatomically couple with posterolateral thalamus (*P L*-*AM*_*c*_) (**Fig. 3E)**.

We then correlated *P L*-*AM*_*c*_ values with *ED*_*pc*1_ loadings (**Fig. 3F)**. We found that cortical parcels with focal thalamic connections preferentially coupled with posterolateral thalamus (*r*_*s*_=-0.57, *p*_*sa*_=0.004) compared to cortical parcels with diffuse thalamic connections, and this relationship was largely consistent across subjects (mean=-0.53, median=- 0.54, SEM=0.004, STD=0.12) (**Fig. 3G**).

We replicated these results using an alternate calculation for the mediolateral thalamic gradient (**Fig. S26**). Additionally, we conducted specificity analyses using the dorsoventral gradient and combinations of the anteroposterior, mediolateral, and dorsoventral gradients (**Fig. S27**). Across subjects, we found that the strongest anatomical coupling association was between *ED*_*pc*1_ loadings and the anteromedial-posterolateral thalamic gradient (**Fig. S27I**).

### The extent of cortical connections within the thalamus is associated with distinct types of information flow

Given previous reports that feed-forward and feed-back cortico-thalamic connectivity vary across the cortical hierarchy (49), we hypothesized that the extent of cortical connections within the thalamus would also correspond to distinct types of information flow in the cortex. Our findings support this hypothesis, demonstrating that cortical parcels with focal thalamic connections are associated with faster, relay-like feed-forward information flow, whereas cortical parcels with diffuse thalamic connections are associated with slower, modulatory feed-back information flow.

Previous studies have characterized two histologically-defined thalamic subpopulations associated with distinct types of information flow. Here, ‘core’ thalamic neurons project focally to middle cortical layers, while ‘matrix’ thalamic neurons project diffusely to superficial cortical layers (5, 66). Additionally, ‘core’ fibers support relay-like, feed-forward information flow, suited for the relaying information, while ‘matrix’ fibers support slower, feed-back information flow, suited for modulatory information processing (5, 66). Recent BOLD-derived functional connectivity work in humans demonstrated that sensory cortical parcels functionally couple with ‘core’ thalamus while associative cortical parcels functionally couple with ‘matrix’ thalamus (64). Based on this evidence, we hypothesized that cortical parcels with more focal thalamic connections would anatomically couple with ‘core’ thalamus, while cortical parcels with more diffuse thalamic connections would anatomically couple with ‘matrix’ thalamus.

To test our hypothesis, we used data from the Allen Human Brain Atlas to estimate the relative mRNA expression levels of calcium-binding proteins Parvalbumin (PVALB), which is more highly expressed in ‘core’ thalamus, and Calbindin (CALB1), which is more highly expressed in ‘matrix’ thalamus. Differences in their mRNA expression index each thalamic voxel’s position along the core-matrix thalamic gradient (*CP*_*t*_) derived from Muller & Munn et al. (64) (**Fig. 4A**). Remarkably, the *CP*_*t*_ gradient strongly correlated with the anteromedial-posterolateral thalamic gradient (*r*_*s*_=0.84) (**Fig. 4B**). While other calcium-binding proteins, like Calretinin, are expressed by thalamic neurons as well (67), we did not consider them in this study.

**Fig. 4.**
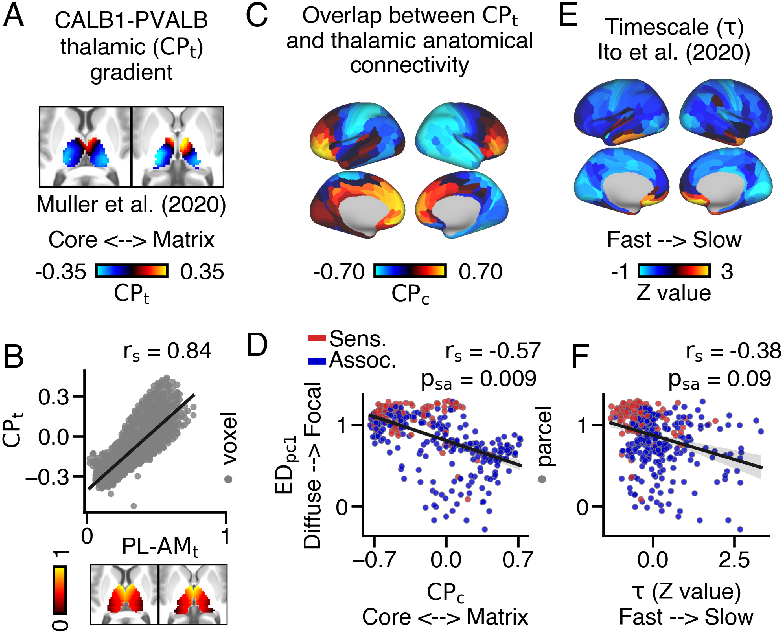
Mapping between cortical variation in the extent of thalamic connectivity patterns, anatomical overlap across thalamic subpopulations, and intrinsic timescale. (**A**) Thalamic gradient reflecting the relative mRNA expression of Parvalbumin (PVALB) and Calbindin (CALB1)-expressing thalamic subpopulations (*CP*_*t*_), which index ‘core’ and ‘matrix’ thalamic subpopulations respectively. (64). Warmer colors reflect higher CALB1 expression relative to PVALB. **(B)** *CP*_*t*_ and *P L*-*AM*_*t*_ values strongly correlated with one another. **(C)** Cortical map reflecting each cortical area’s anatomical overlap across *CP*_*t*_ values (*CP*_*c*_). Warmer colors reflect preferential coupling with CALB1-expressing thalamic populations and cooler colors reflect pref-erential coupling with PVALB-expressing thalamic populations. **(D)** *ED*_*pc*1_ loadings and *CP*_*c*_ values significantly correlated with one another. **(E)** Cortical map of z-transformed intrinsic timescale (*τ*), calculated from group-averaged resting-state data (65). **(F)** *ED*_*pc*1_ loadings and z-transformed *τ* values exhibited a trending correlation when accounting for spatial-autocorrelation.

Next, for each cortical parcel, we correlated their streamline counts within ipsilateral thalamic voxels with those voxels’ *CP*_*t*_ value, resulting in a cortical map of *CP*_*c*_ values (**Fig. 4C**). Here, warmer colors reflect cortical parcels that preferentially target CALB1-expressing ‘matrix’ thalamus while cooler colors reflect cortical parcels that preferentially target PVALB-expressing ‘core’ thalamus.

*ED*_*pc*1_ loadings significantly corresponded with *CP*_*c*_ values (**Fig. 4D**), suggesting that cortical parcels with focal thalamic connections preferentially coupled with ‘core’ thalamus, while cortical parcels with diffuse thalamic connections preferentially coupled with ‘matrix’ thalamus (*r*_*s*_=0.57, *p*_*sa*_=0.009). Moreover, functionally-defined association cortical parcels exhibited higher anatomical *CP*_*c*_ values compared to functionally-defined sensory cortical parcels **(Fig. S28)**.

We next compared the extent of each cortical parcel’s anatomical connections within the thalamus and the intrinsic timescale of their resting-state functional connectivity. Cortical parcels are known to exhibit differences in the timescales of their intrinsic BOLD fluctuations, such that cortical parcels associated with feed-forward processing operate at relatively faster timescales compared to cortical parcels associated with feed-back processing (65, 68). We hypothesized that cortical parcels with more diffuse thalamic connections would exhibit longer intrinsic timescales. To test this, we compared *ED*_*pc*1_ loadings and standardized intrinsic timescale values (*τ*), derived from resting-state functional connectivity from Ito et al. (65) (**Fig. 4E**). We found a modest correlation between *ED*_*pc*1_ loadings and *τ* values, indicating that cortical parcels with focal thalamic connections operated at faster timescales at rest. However, this relationship only showed a trending significance when accounting for spatial auto-correlation (*r*_*s*_=-0.38, *p*_*sa*_=0.09) (**Fig. 4F**).

### Hierarchical variation of the extent of cortical connections within the thalamus is generalized in macaque

Following previous work comparing cortico-thalamic connectivity between humans and macaques (e.g., 41), we compared the extent of anatomical connections within the thalamus between sensory and association cortical areas in macaque monkeys. We hypothesized that cortical variation of the extent of connections within the macaque thalamus would similarly vary along the cortical hierarchy as in humans, albeit to a lesser extent. To test this, we analyzed tractography data from six post-mortem macaque brains, obtained from 7T diffusion MRI scans. We found that the cortical variation in the extent of anatomical connections within the thalamus was generalized in macaques.

We parcellated macaque tractography data to derive connectivity between 128 cortical parcels, derived from the Markov atlas, and ipsilateral thalamic voxels (69). Macaque M1 (area F1; magenta panel) projected to the lateral portion of the thalamus, while DLPFC (area 9/46d; cyan panel) projected to medial and anterior thalamic regions (**Fig. 5A**). Comparing ED values across thresholds, we observed greater similarity between M1 and DLPFC in macaque compared to human. We then performed PCA using the ED matrix to obtain macaque *ED*_*pc*1_ loadings, which accounted for 82% of the variance **(Fig. 5B-C)**.

**Fig. 5.**
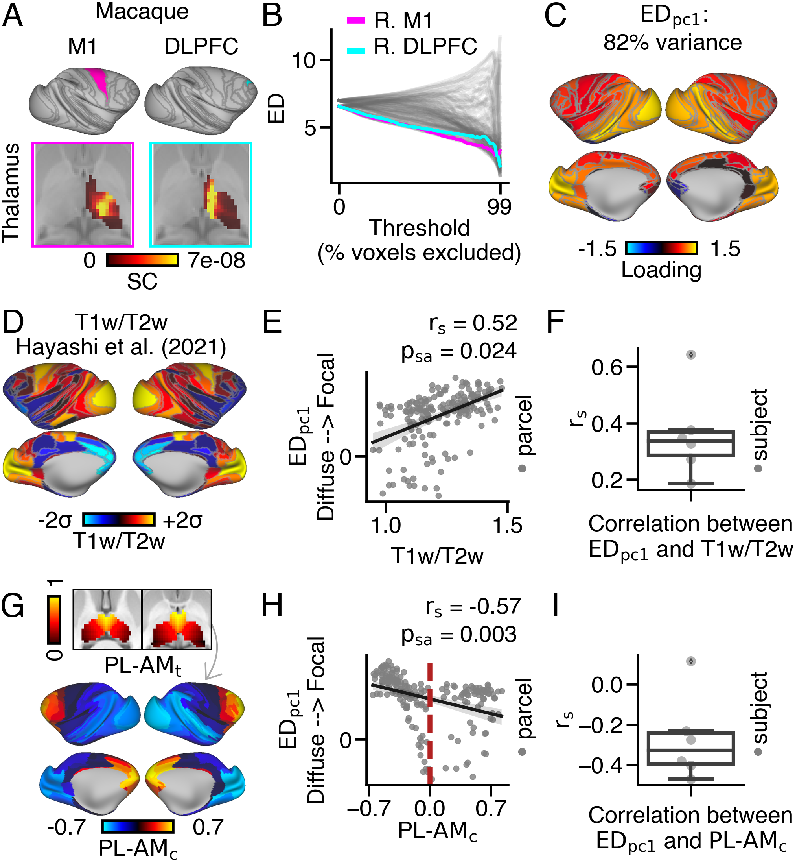
Cortical variation of the extent of anatomical connections within the macaque thalamus. **(A)** Exemplar thalamic connectivity patterns for M1 (area F1; magenta) and DLPFC (area 9/46d; cyan), parcellated using the Markov Atlas (69). **(B)** ED matrix. **(C)** Cortical *ED*_*pc*1_ loading map. **(D)** Cortical myelin map, calculated by averaging T1w/T2w values across 30 macaque monkeys (70). **(E)** Across cortex, *ED*_*pc*1_ loadings positively correlated with T1w/T2w values at the group level and **(F)** at the subject level. **(G)** Cortical map reflecting anatomical coupling with the anteromedial-posterolateral thalamic spatial gradient (*P L*-*AM*_*t*_). Warmer colors reflect cortical parcels that preferentially coupled with anteromedial thalamus (*P L*-*AM*_*c*_). **(H)** Cortical parcels with higher *ED*_*pc*1_ loadings preferentially coupled with posterolateral thalamus. The red dashed line represents parcels with weak preferential coupling the *P L*-*AM*_*t*_ gradient. **(I)** On average, *ED*_*pc*1_ loadings and *P L*-*AM*_*c*_ value negatively correlated across monkeys.

To test if *ED*_*pc*1_ loadings differed between sensory and association cortical parcels in macaques, we correlated group-averaged macaque *ED*_*pc*1_ loadings and group-averaged T1w/T2w values obtained from Hayashi et al. (70) (**Fig. 5D**). We observed a strong positive correlation between *ED*_*pc*1_ loadings and T1w/T2w values at the group level (*r*_*s*_=0.52, *p*_*sa*_=0.024) (**Fig. 5E**), which was consistent on average across subjects (mean=0.36, median=0.34, SEM=0.06, STD=0.15). Surprisingly, no group differences were observed between humans (mean=0.35, STD=.09) and macaques (mean=0.36, STD=.15) (*t*=0.31, *p*=0.75) (**Fig. 5F**; **Fig. S29D**).

We also conducted further analyses to investigate whether cortical parcels with focal thalamic connections preferentially coupled with posterolateral thalamus in macaques. Consistent with the human data, macaque *ED*_*pc*1_ loadings showed a strong negative correlation with *P L*-*AM*_*c*_ values at both the group level (*r*_*s*_=-0.57; *p*_*sa*_=0.003) and the subject-level (mean=-0.27, median=-0.33, SEM=0.086, STD=0.21) **(Fig. 5G-I**). Similar to the human data, the strongest association between *ED*_*pc*1_ loadings and thalamic spatial gradients was with the anteromedial-posterolateral gradient in macaques (**Fig. S30**).

We also compared the correlation between *ED*_*pc*1_ loadings and overlap across each thalamic spatial gradient between species using a two-way ANOVA. We found a significant difference between humans (mean=-0.32, STD=0.25) and macaques (mean=-0.20, STD=0.22) (F(1,8)=24.5, p < 0.001), a significant main effect of gradient (p < 0.001), and no interaction effect (p=0.12). Post-hoc tests showed that only the correlation between *ED*_*pc*1_ loadings and *PL AM*_*c*_ values were significantly different between humans (mean=-0.53, STD=0.12) and macaques (mean=-0.27, STD=0.21) (p=0.041) (**Fig. S29E**).

## Discussion

This study contributes to the rich body of literature investigating the organization of cortico-thalamic systems in human and non-human primates. Prior research has shown that features of thalamocortical connectivity differ between sensory and association systems, and our work advances this understanding by demonstrating that these systems also differ in the pattern and spatial extent of their anatomical connections within the thalamus. Using dMRI-derived tractography across species, we show that these connectivity patterns vary systematically along the cortical hierarchy in both humans and macaques. These findings are critical for establishing the anatomical architecture of how information flows within distinct cortico-thalamic systems. Specifically, we identify reproducible tractography motifs that correspond to sensori-motor and association circuits, which were consistent across individuals and generalize across species. Collectively, this study offers convergent evidence that the spatial pattern of anatomical connections within the thalamus differs between sensory and association cortical areas, which may support distinct computations across cortico-thalamic systems.

### The spatial extent of anatomical connections within the thalamus varies across the cortical hierarchy

Here we replicate findings from prior tracer and tractography studies, providing confirmation that each cortical area exhibits a distinct pattern of anatomical connectivity within the thalamus (16, 18–20, 22, 23, 27–29, 41). Moreover, this study’s findings are consistent with previous animal studies that demonstrate a correspondence between thalamic organization and the sensory-association cortical hierarchy (49–52, 58, 66). Here, we build on this body of work to show, for the first time, that sensory cortical areas project more focally within the thalamus relative to association cortical areas. Our findings suggest that the spatial extent of thalamic connectivity patterns is a key distinguishing feature of cortico-thalamic circuits associated with distinct types of information flow. In line with this notion, other quantitative measures derived from dMRI, including connectivity strength and microstructure, did not significantly vary along the cortical hierarchy.

We also offer convergent evidence supporting the notion of hierarchical anatomical variation by demonstrating that cortical areas with focal thalamic connections preferentially target posterolateral thalamus, associated with the feed-forward relay of sensory information, relative to cortical areas with diffuse thalamic connections connections (58). This is consistent with previous studies showing that the organization of thalamic functional connectivity, microstructure, and gene expression vary along the anteroposterior and mediolateral thalamic axis (16, 21, 59, 71–73). Prior work has shown that axon guidance cues along the anteromedial-posterolateral thalamic gradient shape the topography of thalamo-cortical connections (74). Powell et al. provide insight into the molecular mechanisms that may shape cortex-to-thalamus connectivity patterns. Here, it will be vital to characterize the mechanisms that give rise to hierarchical differences in the spatial extent of anatomical connections within the thalamus, which may be critical for distinct information flow across cortico-thalamic systems and compromised in neurodevelopmental disorders (75–77).

Previous dMRI studies in macaque have identified both similarities and differences in cortico-thalamic anatomical connectivity relative to humans (78). Despite notable interspecies differences in many cortical areas, we found that the extent of connections within the thalamus followed similar patterns of variation across cortex in macaques compared to humans. Such cross-species dMRI analyses provide a bridge with the extensive tract tracing literature in non-human primates. The integration of macaque tractography and tracer data in cortico-thalamic systems remains limited (79, 80), but future studies incorporating tracer data could provide more specific insights into cortico-thalamic system organization by considering directionality, as feed-forward and feed-back thalamic connections vary across cortical areas and may provide more specific insights into the functional roles of these circuits. On the other hand, dMRI-derived tractography data can address gaps in the non-human primate tracer literature. For instance, whole-brain tractography within a single animal is relative easy to acquire. Thus, these data can be used to examine contralateral thalamic connections, which have received less attention in macaque tracer studies (19, 81, 82). Our study shows that cortical parcels with diffuse ipsilateral thalamic connections also have diffuse bilateral thalamic connections, but this relationship wasn’t as strong for contralateral connections. Future studies that examine bilateral and contralateral thalamic connectivity patterns can provide insights the role of contralateral thalamic connections and how cortico-thalamo-cortical circuits influence inter-hemispheric communication.

### Variation in the extent of cortico-thalamic projection patterns and implications for different levels of anatomical architecture

Thalamic and cortical neuronal fibers exhibit distinct patterns in their axonal projections, such that some fibers have axonal projections that project focally while others have more extensive axonal projections (5, 19, 66). Specifically, ‘core’ and ‘matrix’ thalamic neurons exhibit differences in the extent of their axonal terminations, as discussed previously (66). Moreover, prefrontal cortical neurons have been observed to exhibit either dense, focal projections to the ipsilateral thalamus while others have sparse, diffuse projections to bilateral thalamus (19, 83). These differences in the extent of axonal terminations are thought to support different types of neural computations (6), and we hypothesized these distinct patterns of neuronal connectivity may also be reflected in the patterns of large-scale white matter tract terminations within the thalamus.

In this study, we present the first quantitative evidence that the extent of cortical anatomical connections within the thalamus distinguishes cortico-thalamic white matter tracts, mirroring variation seen at the level of cortico-thalamic neurons. Furthermore, we linked human neural gene expression and tractography to show that cortical areas with focal and diffuse thalamic connections preferentially couple with PVALB-expressing ‘core’ and CALB1-expressing ‘matrix’ thalamus, respectively. This finding highlights a conserved anatomical principle of variation between individual neuronal fibers and large-scale white matter tracts within cortico-thalamic systems. This finding suggests that cortico-thalamic white matter tracts may be composed of individual neuronal fibers with similar termination patterns, which can be more precisely tested in animals.

Our findings align with theoretical proposals that hypothesize similar principles governing connectivity at multiple levels of analysis (84, 85). Moreover, it implies that principles of variation at the level of thalamic fibers can provide insights into the properties of large-scale white matter tracts. For instance, thalamic neurons vary in the extent of their axonal projections across cortical layers (5, 66). Furthermore, some thalamic nuclei project to larger swaths of cortex compared to others (27). Future tractography studies can investigate if the extent of thalamic connections within individual cortical areas also varies across the cortical hierarchy.

### The spatial properties of thalamic connectivity patterns provide insight into the role of the thalamus in shaping brain-wide information flow

While early studies suggested that the thalamus primarily relays signals through parallel and segregated circuits. Now, accumulating empirical and computational evidence supports the notion that the thalamus is also a crucial integration hub capable of coordinating and sustaining signals across the cortex (11, 12, 52, 86–88). The structural properties of cortico-thalamic circuits invariably constrain the types of computations these circuits can support (6), and prior work has established a relationship between the extent of axonal terminations and both feed-forward and feed-back information flow in cortico-thalamic systems (19, 66). While the present findings do not directly support directional interpretations, our data raise the possibility that diffuse thalamic connections are associated with slower, feed-back information flow that may support integrative information transmission, whereas focal thalamic connections are associated with faster, feed-forward information flow that may support the relay of sensory information. This hypothesis will likely require further investigation using causal evidence derived from animal studies.

Complementary functional neuroimaging studies have revealed areas of signal integration and segregation within the human thalamus (61, 89, 90). While the lack of recurrent excitatory connections within the thalamus supports segregated information slow, the anatomical basis for information integration within the thalamus is not fully understood. It has been proposed that overlapping terminations within the thalamus may support information integration (23–25, 42). This is supported by work showing that some thalamic neurons receive convergent input from multiple cortical areas (86). In this study, we demonstrate that association cortical areas exhibit diffuse anatomical connections within the thalamus. This may enable these cortical areas to integrate information from distributed areas across the cortex, a critical mechanism supporting higher-order neural computations. Specifically, because thalamocortical connectivity is organized topographically, a cortical area that projects to a larger set of thalamic subregions has the potential to communicate with many other cortical areas. This observation aligns with findings from Phillips et al. that area 24 exhibited the most diffuse anatomical terminations across the mediodorsal nucleus of the thalamus relative to other prefrontal cortical areas (23), which was speculated to support the integration of signals within the prefrontal cortex (42).

Circuits connecting the cortex, thalamus, and the basal ganglia have also been implicated in integrative information transmission (25, 91). The pattern of anatomical connections between the basal ganglia and thalamus may also exhibit variation in their spatial properties. Future studies can examine the spatial properties of anatomical connections between the thalamus and basal ganglia and determine if such variation corresponds with integrated or segregated information flow within cortico-basal ganglia-thalamo-cortical functional interactions.

We also found that ‘first-order’ thalamic nuclei preferentially coupled with cortical parcels with focal thalamic connections. However, our results did not support the hypothesis that ‘higher-order’ thalamic nuclei preferentially couple with cortical areas with diffuse thalamic connections. Instead, these nuclei coupled with cortical areas with both focal and diffuse thalamic connections. Previous studies have shown that ‘higher-order’ thalamic nuclei receive input from cortical and subcortical regions (92, 93). The integration of feed-forward sensory signals and modulatory feed-back signals from cortex could be one mechanism for how the integration of higher-order and first-order signals in the thalamus may support complex cognitive functions (94).

Our findings offer an anatomical framework to complement the findings of a previous investigation on cortico-thalamic functional coupling (64). Muller & Munn et al. (64) examined cortico-thalamic BOLD-derived functional connectivity and demonstrated that sensory cortical areas exhibited stronger functional coupling with ‘core’ thalamus, whereas association cortical areas exhibited stronger functional coupling with ‘matrix’ thalamus, and this dichotomy was found to align with patterns of whole-brain dynamics. However, the specific ways in which thalamo-cortical anatomical constraints may shape functional connectivity remain unknown. A possible future direction would be to investigate how thalamo-cortical anatomical connections contribute to both intra- and inter-hemispheric functional interactions. Lastly, these data imply that cortical areas with diffuse thalamic connections may selectively overlap with thalamic subregions implicated in functional integration and multimodal cognitive processes (61, 90). This observation warrants future investigations in combination with functional modalities.

## Study limitations

While powerful, probabilistic tractography has notable limitations, such as the possibility of producing false positives and lack of directionality information (95, 96). We performed extensive control analyses to identify any inter-subject or inter-cortical confounds. We did not identify any factors that were strongly associated with the *ED*_*pc*1_ measure used to index the extent of thalamic connectivity patterns. We did observe that cortical curvature and sulcal depth were positively correlated with mean streamline count, potentially reflecting gyral bias. This observation should be considered in future tractography studies.

Furthermore, we observed significant individual variation in thalamic connectivity patterns. Notably, we identified biologically implausible tractography patterns in some subjects. The nature of individual differences in thalamic connectivity patterns, whether they represent true individual variation or false positive connections, remains unclear, but it is likely a combination of both. While such variation did not appear to be associated with differences in the extent of thalamic connections between sensory and association cortical areas, our observations are consistent with previous reports demonstrating that tractography data have inherent limitations and should be interpreted with caution (e.g., 95, 96).

We also replicated our findings in macaque monkeys. Macaque dMRI data help mitigate limitations related to lower resolution, shorter collection time, and motion bias in human dMRI studies (44). Tractography-based anatomical connectivity has shown strong agreement with invasive tract tracing studies conducted in monkeys, providing validation for its use (44, 97–103). It is worth noting that the majority of these investigations have primarily focused on cortico-cortical connections and connectivity at the areal level. In comparison, studies directly comparing tractography- and tracer-derived connectivity in thalamo-cortical systems are limited (96, 104). Consequently, future research should prioritize examining the correspondence between tracer and tractography-derived anatomical terminations within subcortical gray matter structures. Furthermore, future studies should investigate factors that may contribute to the presence of biologically implausible connections at the subject level. This is vital for a comprehensive understanding of inter-individual variation of anatomical connectivity and its implications for cognition and behavior in both health and disease.

## Conclusions

The thalamus plays a key role in sensory and association cognitive computations (8, 42, 105). Dysfunction of the thalamus has been linked to severe neuropsychiatric disorders, such as psychosis spectrum disorders (106–111), and the symptoms of these disorders have been associated with abnormal anatomical cortico-thalamic connectivity (75–77). However, our understanding of the role of the thalamus in healthy information transmission and its dysfunction in neuropsychiatric illness has been hindered by limited knowledge of the underlying circuitry, especially in humans (6, 112).

Since the first *in vivo* examinations of cortico-thalamic connectivity in humans (37, 38), neuroimaging studies have made significant progress in mapping thalamo-cortical circuitry and characterizing its role in shaping whole-brain functional interactions (56, 113). This study provides quantitative evidence that the spatial properties of anatomical connections within the thalamus vary across the cortex, following established hierarchical principles of cortical organization, which may reflect variations associated with different types of information flow. Our study highlights that an in-depth investigation of cortico-thalamic anatomical circuitry can offer insights into how the thalamus may support distinct types of information flow throughout the brain, which is critical for the computations that enable higher-order cognition in humans.

### Experimental procedures

#### Human dataset and diffusion processing pipeline

We obtained minimally pre-processed 1.25 mm isotropic 3T dMRI data for 828 healthy adults from the Washington University – Minnesota (WU-Min) Human Connectome Project (HCP). The imaging protocol details can be found at the following link: https://protocols.humanconnectome.org/HCP/3T/imaging-protocols.html (47, 114).

To generate dMRI-derived probabilistic tractography data, we used the Quantitative Neuroimaging Environment & Toolbox (QuNex) (115). Specifically, FSL’s Bedpostx was employed to estimate diffusion parameters, including up to three fiber orientations per voxel, using a model-based deconvolution approach with zeppelins (35, 116, 117). The parameters used were as follows: burn-in period of 3000, 1250 jumps (sampled every 25), automatic relevance determination, and Rician noise. We then obtained whole-brain probabilistic tractography using FSL’s Probtrackx (33, 38, 118). We performed dense gray-ordinate-by-gray-ordinate streamline connectivity, seeding from each white ordinate 3000 times (shown in all figures unless otherwise specified), and from each gray-ordinate 10,000 times **(Fig. S19)** with distance correction. Streamline length data was also extracted. This produced a dense, 91,282 × 91282 grey-ordinate, whole-brain streamline count connectivity matrix.

We then performed several processing steps on the FSL-generated dense tractography data **(Fig. S1)**. To account for inter-subject streamline count differences, dense streamline count data were waytotal normalized. We further applied log normalization to account for distance effects. Each subject’s data were then parcellated along one dimension, by averaging the streamline counts for all grey-ordinates within a cortical parcel. The parcellation used was defined by Ji et al. (48), and the analysis was restricted to the 360 symmetrical bilateral cortical parcels defined by Glasser et al. (45). These data were masked with the thalamic gray matter mask used during tractography, which consisted of 2539 voxels **(Fig. S21)**. Group-level cortical-parcel by thalamic-voxel streamline count connectivity matrices were generated by averaging data across participants. In some visualizations, standardized streamline counts were shown, which were z-scored for each cortical parcel. Finally, before group averaging, an alternative processing step was performed which consisted of regressing cortical curvature from streamline counts for each subject. The residuals from this regression were then group-averaged **(Fig. S1, step 5)** and used for supplementary analyses.

We replicated our findings using multiple functionally and structurally-defined thalamic masks derived from different atlases: the Yeo 2011 parcellation (119) (https://github.com/ryraut/thalamic-parcellation), Melbourne Atlas (120), and the Morel thalamic atlas (57) **(Fig. S21)**. Furthermore, we replicated the main results using dense ED values **(Fig. S20)** and an alternative seeding strategy, which consisted of seeding each grey-ordinate 10,000 times **(Fig. S19)**. We also found no difference after regressing total streamline count to ipsilateral cortex **(Fig. S31)**.

#### Human BOLD acquisition and processing

For each subject, we obtained four runs of minimally preprocessed blood-oxygen-level-dependent (BOLD) resting-state data from the HCP (Atlas_MSMAll_hp2000_clean). The first 100 frames of each BOLD time series were dropped, and the data were demeaned. The four resting-state scans were concatenated in the order of 2-1-4-3. These data were parcellated using the parcellation defined by Ji et al. (48), and we included only the 360 cortical parcels defined by Glasser et al. (45). Functional connectivity was estimated using pairwise Pearson correlations between the time series of each cortical parcel, resulting in a parcellated functional connectivity matrix. This matrix was then used as input into a principal component analysis (PCA) to derive the first and second principal functional gradients. See *Deriving cortical gradients* for more details.

#### Macaque dataset and diffusion processing pipeline

We obtained diffusion-weighted 7T MRI data with 0.6 mm isotropic resolution from a previously collected dataset of six postmortem macaques (4-16 years old), as described in previous work (121, 122). These data are publicly available through the PRIMatE Data Exchange (PRIME-DE) repository (http://fcon_1000.projects.nitrc.org/indi/PRIME/oxford2.html) (123). Nonlinear surface transformation to macaque F99 standard space was performed, as described elsewhere (124, 125). Subcortical structures were registered to F99 standard space using FNIRT and templates from the HCP Non-Human Primate Minimal Preprocessing Pipelines (47, 70).

Streamline count connectivity matrices for the macaques were derived using FSL’s Bedpostx and Probtrackx pipeline. We seeded each white-ordinate 3,000 times to obtain a gray-ordinate-by-gray-ordinate connectivity matrix. The parameters used were the same as those used for tractography in the human data, with the exception that no distance correction was applied and the step length was reduced from 0.5 to 0.2. The macaque data were waytotal and log normalized, and then parcellated using the Markov 2014 atlas (69, 126). This workflow resulted in a parcel-by-dense (128×71401) connectivity matrix for each macaque. These data were masked with the thalamic mask used for tractography seeding, which consisted of 1539 voxels (649 right; 890 left), and data were then averaged to create a group matrix.

#### Framework to quantify the extent of thalamic connectivity patterns via Euclidean distance (ED)

We used Euclidean distance (ED) to quantify the extent of each cortical area’s thalamic connectivity patterns. Probabilistic tractography data require thresholding before the ED calculation. To avoid the selection of an arbitrary threshold (34, 36), we calculated ED for a range of thresholds **(Fig. 1A)**. Our thresholding framework uses a tractography-derived connectivity matrix as input. We iteratively excluded voxels with lower streamline counts for each cortical parcel such that the same number of voxels was included at each threshold. At each threshold, ED was calculated between the top x% of thalamic voxels with the highest streamline counts. This produced a matrix of ED values (360 cortical parcels by 100 thresholds). This matrix was used as input into a PCA to derive a single loading for each cortical parcel. While alternative thresholding approaches have been proposed, this framework optimizes the examination of spatial patterns by proportionally thresholding the data, enabling equitable sampling of each cortical parcel’s streamline counts within the thalamus.

This approach controlled for inter-areal differences in anatomical connection strength that could confound the ED estimates. In contrast, a global threshold, which is applied to all cortical areas, may exclude all thalamic streamline counts for some cortical areas that are more difficult to reconstruct, thus making it impossible to calculate ED for that cortical area, as there are no surviving thalamic voxels from which to calculate ED. This would be especially problematic for white matter tracts that are more difficult to reconstruct (e.g. the auditory radiation), and cortical areas connected to the thalamus by those white matter tracts would have a disproportionate number of thalamic voxels excluded when using a global threshold.

We iteratively thresholded tractography-derived cortico-thalamic connectivity data from 828 healthy adults from the HCP by excluding 0% to 99% of thalamic voxels with the lowest streamline counts for each cortical parcel **(Fig. 1B)**. Pairwise ED was calculated between thalamic voxels that survived thresholding for 100 thresholds, which resulted in an ED value for each threshold for each cortical parcel (Fig. 1C). The pairwise ED calculation was calculated using the following equation:

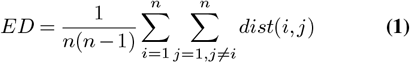

Where:

- *ED* is the average pairwise ED,
- *n* is the number of surviving thalamic voxels,
- *i* and *j* are indices of thalamic voxels,
- dist(*i,j*) is the ED between thalamic voxels *i* and *j*.

The Euclidean distance between two thalamic voxels, dist(*i,j*), was calculated using the equation:

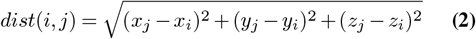

where *x*_*i*_, *y*_*i*_, *z*_*i*_ and *x*_*j*_, *y*_*j*_, *z*_*j*_ represent the coordinates of the thalamic voxels *i* and *j* in 2mm space. This calculation was performed for each cortical parcel across its surviving thalamic voxels (i.e., the top x% of thalamic voxels with the highest streamline counts) for each threshold, generating a cortical parcel-by-threshold matrix of ED values. The procedure was performed separately for the left and right thalamus. In the case of bilateral connectivity patterns, the ED values for each threshold were summed across the left and right thalamus. All analyses used ipsilateral ED values, unless otherwise specifies in supplementary analyses.

Additionally, we calculated relative ED (rED) to account for size differences between left and right thalamus:

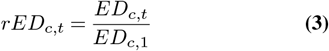

Where:

- *rED* denotes relative ED,
- *ED*_*c,t*_ is the ED value for each cortical parcel *c* at threshold *t*,
- *ED*_*c*,1_ is the ED value for each cortical parcel *c* at threshold 1 (when no threshold is applied and no voxels are excluded).

In this study, due to the similarity in size between the left and right thalamus, *ED*_*pc*1_ and *rED*_*pc*1_ loadings were nearly identical (Fig. S14).

#### Measure calculations

##### *ED*_*pc*1_ loading calculation

PCA using Singular Value Decomposition was performed using the threshold by cortical parcel (100×360) ED matrix. The first principal component (*ED*_*pc*1_) is a vector of the 360 loadings from PC1. The loadings were calculated following the equation:

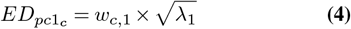

Where:

- *w*_*c*,1_ is the first PC’s eigenvector for cortical parcel c,
- *λ*_1_ is the first PC’s eigenvalue.

This procedure was also performed for group-averaged dense cortico-thalamic connectivity data to examine the extent of anatomical connections within the thalamus for 59,412 cortical vertices **(Fig. S20)**.

##### *ED σ* calculation

*ED σ* is the population standard deviation of ED across thresholds for each cortical parcel. It was calculated following the equation:

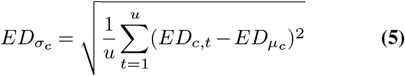

Where:

- 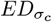 is the standard deviation of ED values across thresholds for cortical parcel *c*,
- *u* is the total number of thresholds (in this case 100),
- *ED*_*c,t*_ is the ED of surviving thalamic voxels for parcel *c* at threshold *t*,
- 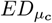is the mean ED for cortical parcel *c* across *u* thresholds.

In addition to the standard deviation of ED across all thresholds, we also calculated the standard deviation (STD) of ED across specific ranges of thresholds (Fig. S6). We then correlated these values with *ED*_*pc*1_, T1w/T2w, and *RSFC*_*pc*1_ measures.

##### Mean SC calculation and analysis

The mean streamline count (Mean SC) within the thalamus was calculated for each cortical parcel following the equation:

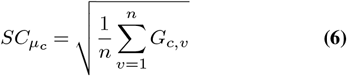

Where:

- 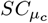 is the square root normalized mean of streamline counts for cortical parcel *c*,
- *v* is the index of the thalamic voxel,
- *n* is the number of thalamic voxels,
- *G*_*c,v*_ is the streamline count between cortical parcel *c* and thalamic voxel *v*.

We then correlated Mean SC with *ED*_*pc*1_, T1w/T2w, *RSFC*_*pc*1_, *RSFC*_*pc*2_, and average streamline length values.

##### Streamline length calculation and analysis

The average streamline length (l) for each cortical parcel was calculated following the equation:

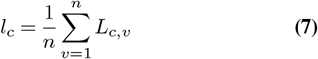

Where:

- *l*_*c*_ is the average streamline length for cortical parcel *c* across thalamic voxels,
- *v* is the index of the thalamic voxel,
- *n* is the number of thalamic voxels,
- *L*_*c,v*_ is the average length of streamlines between cortical parcel *c* and thalamic voxel *v*.

Streamline lengths were obtained from FSL’s probtrackx using the –ompl flag.

##### Isotropy (*I*_*pc*1_) calculation

To quantify the isotropy of thalamic connectivity patterns for each cortical parcel, we modified the thresholding framework shown in **Fig. 1A** and replaced the ED calculated with the following the equation:

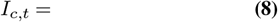

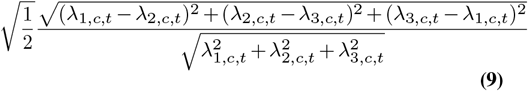

Here, λ values were obtained from the covariance matrix derived from the x, y, and z coordinates of surviving thalamic voxels for each threshold *t* and cortical parcel *c*. The *I*_*pc*1_ measure reflect how evenly spread out each cortical parcel’s anatomical connections are within the thalamus.

This covariance matrix was used as input into a PCA, and the loadings from the first principal component (PC1) were calculated as described in the *ED loading calculation* section.

##### *ED*_*pc*1_ score calculation

PCA was performed on the cortical parcel by threshold (360×100) ED matrix (i.e., the transposed matrix). PC1 scores for each cortical parcel were calculated following the equation:

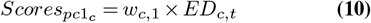

Where:

- 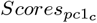 is the PC1 score for cortical parcel *c*,
- *ED*_*c,t*_ is the cortical parcel by threshold ED matrix,
- *w*_*c*,1_ is the eigenvector corresponding to the first PC for cortical parcel *c*.

Mean ED calculation:

The average ED across thresholds was calculated for each cortical parcel by taking the arithmetic mean of the ED values across thresholds. The calculation was performed using the following equation:

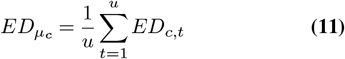

Where:

- 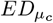 is the mean ED for cortical parcel *c* across *u* thresholds,
- *u* is to the total number of thresholds (in this study we used 100),
- *t* is the index of the threshold,
- *ED*_*c,t*_ is the ED of surviving thalamic voxels for cortical parcel *c* at threshold *t*.

In addition to the mean ED across all thresholds, we also calculated the mean ED across ranges of thresholds (Fig. S6) and compared these values to *ED*_*pc*1_, T1w/T2w, and *RSFC*_*pc*1_ values.

##### Weighted Mean ED calculation

We weighted ED values for each threshold to derive a weighted mean of ED values across thresholds, with more conservative thresholds having higher weights. The calculation was performed using the following equations:

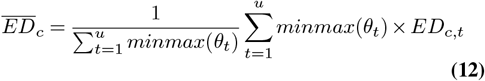

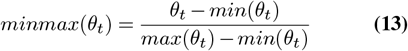

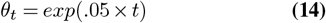

Where:

- 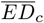is the weighted mean ED for cortical parcel *c* across *u* thresholds,
- *u* is the total number of thresholds (in this study we used 100),
- *t* is the index of the threshold,
- _*θ t*_ is the weight for the *t*^*th*^ threshold.

The weight assigned to each threshold was determined by an exponential function. Specifically, a threshold of *t* = 1, where no voxels were excluded, had a weight of zero, while a threshold of *t* = 100, where 99% of voxels were excluded, had a weight of 1. In other words, less conservative thresholds are ‘discounted’ and contribute less to the overall mean.

##### Streamline count (SC) Skewness calculation

The robust skewness of streamline counts within the thalamus was calculated for each cortical parcel using the mean-median difference standardized by the absolute deviation. The calculation was performed using the following equation:

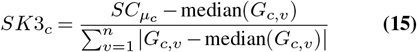

Where:

- *SK*3_*c*_ is the robust skewness of SCs for cortical parcel *c*,
- 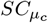is the mean of square-root normalized streamline counts for cortical parcel *c*,
- median(*G*_*c,v*_) is the median of streamline counts between cortical parcel *c* and thalamic voxel *v*,
- *v* is the index of the thalamic voxel,
- *n* is the number of thalamic voxels.

This measure was calculated based on the hypothesis that cortical areas with more focal connections within the thalamus would have a more skewed distribution of streamline counts within the thalamus. However, because skewness is sensitive to outliers, we used a robust measure to discount extreme values.

##### Weighted Nuclei Mean SC calculation

The mean streamline count between each cortical parcel and each thalamic nucleus was weighted by the volume of the nucleus. These weighted values were then summed across thalamic nuclei. The calculation was performed using the following equation:

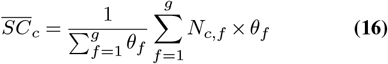

Where:

- 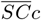is the standardized mean streamline count across all thalamic nuclei, weighted by the volume of each nucleus (*θ*_*f*_),
- *f* is the index of the thalamic nucleus,
- *g* is the total number of thalamic nuclei from the Morel thalamic atlas (g = 28),
- *N c, f* is the mean streamline count between cortical parcel *c* and thalamic nucleus *f*, standardized within cortical parcels.

This measure was calculated based on the rationale that cortical parcels with more diffuse connections within the thalamus would be strongly connected to more thalamic nuclei than those with focal thalamic connections. Because thalamic nuclei have very different sizes, we created weighted the mean streamline counts of thalamic nuclei by their volume. This measure may also be used with parcellated data, when dense data are not available.

### Control analyses

Cortical maps for cortical surface features, including cortical thickness, curvature, sulcal depth, bias field, edge distortion, spherical distortion, and areal distortion, were obtained from the HCP and group-averaged across subjects. The surface area of each cortical parcel from the Glasser et al., atlas was calculated as the number of vertices within each parcel. These cortical maps were then examined for associations with *ED*_*pc*1_ loadings **(Fig. S13)**.

Additionally, fractional anisotropy (FA) and mean diffusivity (MD) values were extracted for each subject using FSL’s DTIFIT. The correlation between the streamline counts of each cortical parcel within the thalamus and their corresponding thalamic FA and MD values was then calculated to index anatomical overlap. These correlations were then correlated with *ED*_*pc*1_ loadings **(Fig. S12)**.

#### Network analysis

Network assignments for the 360 bilateral cortical parcels were derived from by Ji et al. (48). Cortical parcels were categorized into 12 functionally-defined networks, including four sensory networks (somatomotor, SMN; visual 1, VIS1; visual 2, VIS2; auditory, AUD) and eight association networks (cingulo-opercular, CON; default-mode, DMN; dorsal attention, DAN; frontoparietal network, FPN; language, LAN; posterior multimodal, PMM; ventral multimodal, VMM; orbito-affective, ORA) **(Fig. 2A)**. *ED*_*pc*1_ loadings were averaged within these 12 networks and within the sensory and association networks for each subject **(Fig. 2B-C)**. The *ED*_*pc*1_ loadings were averaged within sensory and association networks for each subject and compared using a Wilcoxon signed-rank test **(Fig. 2C)**.

#### Cortical gradients

To capture systematic variation across the cortex, we calculated multiple cortical gradients and compared them to the cortical *ED*_*pc*1_ loading map.

T1w/T2w maps, reflecting cortical myelin content (127), were obtained from the HCP dataset and group-averaged across 828 subjects. Similarly, group-averaged dense T1w/T2w maps from 30 macaques were acquired (https://balsa.wustl.edu/study/Klr0B) (70).

Functional cortical gradients were derived from group-averaged and individual-level (n=828) resting-state functional cortical connectivity matrices. After thresholding to include the top 20% of connections, PCA was performed, and the loadings from the first (*RSFC*_*pc*1_) and second (*RSFC*_*pc*2_) principal components were calculated, following previous work (55, 128).

T1w/T2w values and *RSFC*_*pc*1_ loadings serve as quantitative indexes of cortical hierarchy, with sensory cortical parcels exhibiting higher T1w/T2w values and *RSFC*_*pc*1_ loadings compared to association cortical parcels. Individual-level T1w/T2w maps and *RSFC*_*pc*1_ maps highly correlated with group-averaged maps across subjects (**Fig. S17**). *RSFC*_*pc*2_ loadings index the sensory-association-motor cortical hierarchy, which reflects functional specialization (55).

To examine the anteroposterior cortical gradient, we calculated the Cartesian distance between each cortical parcel and ipsilateral V1, separately for human and macaque data.

#### Non-parametric method for assessing the correspondence of cortical brain maps

We assessed the correspondence between cortical brain maps using Spearman correlations and determined the significance of each correlation using a non-parametric approach. To preserve spatial autocorrelation, we generated 1000 surrogate maps for each target brain map (e.g., T1w/T2w values, *RSFC*_*pc*1_ loadings) using brainSMASH with default parameters (46). Specifically, 500 surrogate maps were generated for the left cortex and 500 for the right cortex. These surrogates were then mirrored contralaterally to create bilateral surrogate maps. We correlated these bilateral surrogate maps with the empirical cortical map (e.g., *ED*_*pc*1_ loadings) to generate a null distribution. Non-parametric p-values were calculated by dividing the number of surrogate maps with a higher correlation coefficient than the empirical value by the total number of surrogates generated (i.e., 1000).

For dense maps, surrogates were generated separately for the left and right cortex using resampling. We used default parameters with the exception of the knn parameter, which was set to the number of vertices in the brain map, following (129).

#### Thalamic nuclei segmentation

Thalamic nuclei were defined using the Morel histological atlas (57), resampled to 2mm Montreal Neurological Institute (MNI) space and converted to cifti format **(Fig. 3A)**. Each thalamic voxel was assigned to one of the 28 thalamic nuclei based on the highest scaled mask value. The sPF nucleus from the Morel atlas was combined with the PF nucleus as their voxels overlapped in 2mm space. Thalamic nuclei were categorized into posterior, medial, lateral, and anterior subdivisions (130), and some were classified as higher-order or first-order (15) and primary, secondary, and tertiary (59) based on prior work. The complete list of 28 thalamic nuclei, along with their abbreviations, is provided in **Table. 1**. The volume of each nucleus was also calculated by counting the number of 2mm thalamic voxels assigned to it **Table. 1**.

For each cortical parcel, we calculated the mean streamline count across thalamic voxels within each thalamic nucleus. This produced a cortical parcel by thalamic nucleus (360×28) streamline count connectivity matrix for each subject. These matrices were then averaged across all subjects and z-scored to visualize the cortical connectivity patterns within each thalamic nucleus **(Fig. 3B)**. The streamline count connectivity matrix was sorted along the y-axis such that cortical parcels with lower average *ED*_*pc*1_ loadings were located at the bottom.

Next, each subject’s *ED*_*pc*1_ loadings were correlated with the Mean SC values across cortex for each of 28 thalamic nuclei, resulting in 28 rho values per subject **(Fig. 3C)**.A positive rho value indicated that a thalamic nucleus preferentially coupled with cortical parcels with focal thalamic connections, while a negative rho value indicated that a thalamic nucleus preferentially coupled with cortical parcels with diffuse thalamic connections. A rho value of zero indicated that a thalamic nucleus had no preference and equally coupled to cortical parcels with focal and diffuse thalamic connections.

To test for differences between subject-level rho values, we conducted a Friedman test to compare the averaged rho values between *ED*_*pc*1_ and Mean SC between posterior, lateral, anterior, and medial thalamic nuclei. Post-hoc analyses were performed using the Wilcoxon–Nemenyi–McDonald–Thompson test, which corrected for family-wise error **(Fig. 3C)**. A similar procedure was conducted to examine the differences in average subject-level rho values between *ED*_*pc*1_ and Mean SC for primary, secondary, and tertiary nuclei (59) **(Fig. S24)**. Additionally, a two-sided Wilcoxon signed-rank test was employed to determine the significance of differences between subject-level rho values for first-order and higher-order thalamic nuclei **(Fig. 3D)**. We also examined correlations between cortical Mean SCs for each thalamic nucleus and the minor and major subdivisions of thalamic nuclei (**Fig. S23**); however, no comparisons survived correction for multiple comparisons using the Holm-Bonferroni correction.

#### Individual variability

Based on tract tracing findings in monkeys, we expected V1 thalamic connections to terminate in visual thalamic areas such as the lateral geniculate nucleus and pulvinar in the posterior thalamus (131). While some subjects exhibited V1 terminations in the visual thalamus, as exemplified by subject 1, others exhibited termination patterns that spread along the anterior and dorsal axis of the thalamus, as seen in subject 2 **(Fig. S18A)**. To quantify these differences, we calculated the ratio of Mean SCs in the visual thalamus relative to Mean SCs outside of the visual thalamus for the right V1 for each subject. Here, a more positive ratio indicated a larger proportion of V1 streamlines terminating in the visual thalamus, which comprised of the pulvinar and lateral geniculate nucleus **(Fig. S18B)**. All subjects had right V1 terminations that preferentially coupled with the pulvinar and lateral geniculate nucleus (ratios > 1). However, many subjects had ratios close to 1, indicating a substantial portion of streamlines terminating outside of the visual thalamus. A strong negative correlation was observed between each subject’s right V1 *ED*_*pc*1_ loading and their right V1 SC ratio, demonstrating that subjects with more diffuse right V1 connections within the thalamus had streamlines that terminated outside the visual thalamus (*r*_*s*_ = -0.81) **(Fig. S18B)**.

Finally, we investigated the hypothesis that V1 terminations outside of visual thalamus would diminish the correspondence between *ED*_*pc*1_ and the T1w/T2w and *RSFC*_*pc*1_ cortical maps. To assess this hypothesis specifically for right V1, we compared each subject’s ratio of streamline counts between visual and non-visual thalamus to their respective Spearman rho values between their *ED*_*pc*1_ loadings and cortical myelin (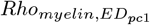; purple) and the principal functional gradient (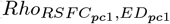; green) **(Fig. S18C)**. We observed moderate positive correlation between these variables, indicating that subjects with weaker correlations between *ED*_*pc*1_ loadings and T1w/T2w and *RSFC*_*pc*1_ values also had more right V1 terminations outside of the visual thalamus. This finding suggests that biologically-unlikely V1 connections that extend outside of visual thalamus may weaken the relationship between *ED*_*pc*1_ loadings and T1w/T2w and *RSFC*_*pc*1_ values. It is important to note that this observation pertains to a single cortical parcel, and further research is needed to elucidate how such biologically-plausible connections may bias the connectivity patterns of other cortical parcels.

#### Cartesian thalamic gradient calculation and analysis

To capture thalamic anatomical coupling along continuous thalamic gradients, we examined the overlap between each cortical parcel’s thalamic streamline counts along Cartesian spatial gradients. We correlated each voxel’s streamline count and position along each thalamic spatial gradient for each cortical area. Specifically, the anteroposterior and dorsoventral gradients were defined by the y and z coordinates (*y*_*t*_ and *z*_*t*_), respectively. The mediolateral gradient was calculated two ways. First, we calculated the distance to the cortical midline (*M*_*t*_) using the formula:

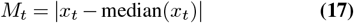

Where:

- *M*_*t*_ is the distance to the cortical midline for thalamic voxel *t*,
- *x*_*t*_ is the x coordinate of each thalamic voxel,
- median(*x*_*t*_) is the median x coordinate across all thalamic voxels.

Next, we calculated the distance to the thalamic midline (*ML*_*t*_) following the equation:

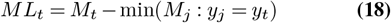

Where:

- *ML*_*t*_ is the distance to the thalamic midline for thalamic voxel *t*,
- *M*_*t*_ is the distance to the cortical midline for thalamic voxel *t*,
- median(*x*_*t*_) is the median x coordinate across all thalamic voxels.
- min(*M*_*j*_ : *y*_*j*_ = *y*_*t*_) is the distance between the cortical midline and the most medial part of the thalamus corresponding to the thalamic voxel’s y-axis position,
- *y*_*t*_ is the y coordinate of a given thalamic voxel,
- *j* represent the indices of thalamic voxels that share the same y coordinate as the thalamic voxel at position *t*.

By subtracting the distance to the midline thalamus from each thalamic voxel’s distance to the cortical midline, we obtained the mediolateral thalamic gradient (see **Fig. S27** to view thalamic gradients).

Since the thalamus sits slightly oblique to the cortical midline, the distance to the midline (*M*_*t*_) exhibited a stronger correspondence with the anteroposterior thalamic gradient compared to distance to midline thalamus (*ML*_*t*_) **(Fig. S26A)**. This demonstrates that the using distance to the cortical midline (*M*_*t*_) does not capture a true thalamic medio-lateral gradient, because it is confounded by the anteroposterior axis. Therefore, we used the distance to midline thalamus (*ML*_*t*_) to quantify the mediolateral thalamic spatial gradient in the main text. Additionally, we replicated our results using a separate calculation for the mediolateral thalamic gradient using the distance to the cortical midline (i.e., using *M*_*t*_ instead of *ML*_*t*_). These findings largely mirrored the main text results, with the anteromedial-posterolateral thalamic gradient showing the strongest relationship with *ED*_*pc*1_ loadings **(Fig. S26)**. These data highlight the need for the careful consideration of which mediolateral gradient may be more appropriate to address hypotheses regarding thalamic organization.

We also examined combinations of the anteromedial, posterolateral, and dorsoventral thalamic gradients. First, we min-max transformed the *y*_*t*_, *z*_*t*_, and *ML*_*t*_ values to scale them between 0 and 1. Then, we calculated the position of each thalamic voxel’s position along six thalamic spatial gradients following the equations:

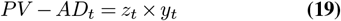

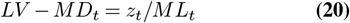

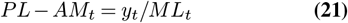

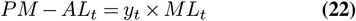

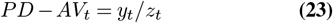

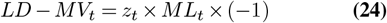

Where:

- *t* denotes index of the thalamic voxel,
- *ML*_*t*_, *y*_*t*_, and *z*_*t*_ denote each thalamic voxel’s position along the mediolateral, anteroposterior, and dorsoventral thalamic gradients,
- *PV* -*AD*_*t*_ : position of thalamic voxel *t* along the anterodorsal-posteroventral gradient,
- *LV* -*M D*_*t*_ : position of thalamic voxel *t* along the mediodorsal-lateroventral gradient,
- *P L*-*AM*_*t*_ : position of thalamic voxel *t* along the anteromedial-posterolateral gradient,
- *PM* -*AL*_*t*_ : position of thalamic voxel *t* along the anterolateral-posteromedial gradient,
- *PD*-*AV*_*t*_ : position of thalamic voxel *t* along the anteroventral-posterodorsal gradient,
- *LD*-*M V*_*t*_ : position of thalamic voxel *t* along the medioventral-laterodorsal gradient.

These values were also min-max scaled between 0 and 1 (see **Fig. S27** to view the thalamic gradients). Using the same overlap procedure described earlier, we generated six cortical maps reflecting anatomical overlap with each thalamic gradient and compared these cortical maps to the empirical *ED*_*pc*1_ cortical map **(Fig. S27)**. We performed this procedure for both humans and macaques (see **Fig. S30** for the macaque gradients).

#### Cross-species comparisons

The *ED*_*pc*1_ calculation framework was applied to 7T dMRI data for six post-mortem macaque monkeys (**Fig. 5A**). ED values were calculated for 100 thresholds and PCA was applied to the resulting ED matrix to extract *ED*_*pc*1_ loadings for 128 cortical areas defined using the Markov atlas (69) (**Fig. 5B-C**).

The relationship between macaque *ED*_*pc*1_ and *ED*_*σ*_ was slightly weaker and more nonlinear compared to the human data, likely due to cortical parcels with exceptionally high ED values, resulting in an underestimation of the extent of connections within the thalamus by the *ED*_*σ*_ measure. Conversely, the *ED*_*pc*1_ measure accurately captured the extent of connections within the thalamus for these cortical parcels (**Fig. S29A-C**).

To determine if *ED*_*pc*1_ loadings differed between sensory and association cortical parcels we used a group-averaged T1w/T2w macaque cortical map derived from data from Hayashi et al. (70) (**Fig. 5D**) and correlated these values with *ED*_*pc*1_ loadings at both the group and subject level (**Fig. 5E-F**). We also performed a two-sample t-test to compare these correlations between species (**Fig. S29D**)

The median correlation coefficient (rho) between ipsilateral *ED*_*pc*1_ loadings and overlap across thalamic spatial gradient was compared between humans and macaques using a 2-way analysis of variance (ANOVA). We determiend if the residuals of the ANOVA model were normally distributed using a QQ plot. Both species exhibited equal variances (Bartlett’s test; p = 0.21), but the gradients, despite having the same sample size, showed unequal variances (Bartlett’s test; p < 0.001). To avoid introducing an artificial interaction between gradients and species, the sign of the rho values between *ED*_*pc*1_ loadings and overlap across the mediolateral gradient was flipped prior to conducting the statistical tests. Post-hoc Tukey’s Honestly Significant Difference (HSD) test for multiple comparisons was then performed (**Fig. S29E**), which showed a species difference only for the anteromedial-posterolateral thalamic gradient,

#### PVALB and CALB1 thalamic gradient calculation and analysis

The thalamic gradient of the relative mRNA levels of Calbindin (CALB1) and Parvalbumin (PVALB) (*CP*_*t*_) was downloaded from https://github.com/macshine/corematrix (64). This map was originally obtained from the Allen Brain Atlas using two probes to estimate PVALB expression (CUST_11451_PI416261804 and (A_23_P17844) and three probes to estimate CALB1 expression (CUST_140_PI416408490, CUST_16773_PI416261804 and A_23_P43197). Additional details are described elsewhere (64). Cooler colors on the *CP*_*t*_ gradient reflect higher relative expression of PVALB, associated with the ‘core’ thalamus, while warmer colors reflect higher relative expression of CALB1, associated with the ‘matrix’ thalamus (**Fig. 4A**). We also correlated the *CP*_*t*_ gradient with the anteromedial thalamic gradient (*P L*-*AM*_*t*_) **(Fig. 4B)**.

To calculate the spatial overlap between each cortical parcel’s thalamic connectivity patterns and ‘core’-’matrix’ thalamic subpopulations, the *CP*_*t*_ map was correlated with each cortical parcel’s thalamic streamline counts. Only voxels with *CP*_*t*_ values that overlapped with the thalamic mask used for tractography were included in these analyses (1,684 bilateral thalamic voxels). This overlap procedure produced a cortical map (*CP*_*c*_) with a single *r*_*s*_ value for each cortical parcel, where cooler values reflect cortical parcels whose thalamic connectivity patterns overlap more with PVALB-expressing thalamic subpopulations, and warmer colors reflect cortical parcels whose thalamic connectivity patterns overlap more with CALB1-expressing thalamic subpopulations **(Fig. 4C)**. Next, we correlated the *CP*_*c*_ cortical map with the *ED*_*pc*1_ cortical map **(Fig. 4D)**. We also correlated *CP*_*c*_ cortical map with the T1w/T2w and *RSFC*_*pc*1_ cortical maps **(Fig. S28B-C)**. We also correlated the *CP*_*c*_ left and right cortical maps with left and right cortical maps of the *ED*_*pc*1_ measure, as the *CP*_*t*_ thalamic map showed an asymmetric topography of PVALB-CALB1 expression **(Fig. S28D)**.

#### Intrinsic timescale

We performed an auto-correlation analysis using intrinsic timescale from BOLD functional connectivity data from the HCP (132). The data were obtained from Ito et al. (2020) (https://github.com/ColeLab/hierarchy2020) (65). For each cortical parcel, an exponential decay function was fit for each HCP subject using the equation:

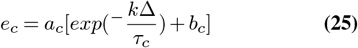

Where:

- *e*_*c*_ is the exponential decay function fit for each cortical parcel *c*,
- *a*_*c*_ is a scaling factor,
- *b*_*c*_ is an offset,
- _*τ c*_ reflects the rate of decay (i.e., intrinsic timescale),
- *k* Δ is the time lag (we used a lag of 100 timepoints).

The model was fit individually for each cortical parcel using the ‘Trust Region Reflective’ algorithm (scipy.optimize.curvefit), and the data were standardized. These *τ* values were then correlated with *ED*_*pc*1_ loadings.

#### Visualization

All cortical brainmaps were generated using Connectome Workbench. Subcortical axial visualization were generated using Nilearn Plotting in python. Figures were constructed in python with CanD v0.0.2 (https://github.com/mwshinn/CanD).

## Data and Code Availability

The neuroimaging files used in this study will be made available on the Brain Analysis Library of Spatial maps and Atlases (BALSA) website. The 1mm Morel thalamic atlas was obtained elsewhere with permission (57). As such, we do not provide the thalamic nuclei files directly, but we will provide the code to create the labels for Morel nuclei in 2mm space. All code related to this study will be made publicly available on BitBucket. All analyses were implemented using Python (version 3.10) and the following packages were used: numpy v1.21.2 (133), pandas v1.4.4 (134), scipy v1.10.1 (135), nibabel v3.2.1 (https://zenodo.org/record/7795644), seaborn v0.12.2 (136), sklearn v0.10.0 (137), matplotlib v3.4.3 (138), wbplot (https://github.com/jbburt/wbplot), pingouin v0.5.3 (https://pingouin-stats.org/build/html/index.html), statsmodels v0.13.1 (139), and nilearn (140).

## ACKNOWLEDGEMENTS

We thank Jie Lisa Ji, Grega Repovš, Jure Demsar, and Takuya Ito for assistance with data processing and analysis and Amy F.T. Arnsten, Jane R. Taylor, John H. Krystal, Maxwell Shinn, Rachel Cooper, Jacob A. Miller, and Warren W. Pettine for helpful discussions. This work was supported by the Gruber Foundation (AMH) and the National Institute of Mental Health (Grant Nos. U01 MH121766-03, R01 MH112189) (AA). Diffusion data collection and sharing for this project was provided by the Human Connectome Project (HCP) and the PRIMatE Data Exchange (PRIME-DE).

