## Supplemental_Figures for "The spatial extent of anatomical connections within the thalamus varies across the cortical hierarchy in humans and macaques"

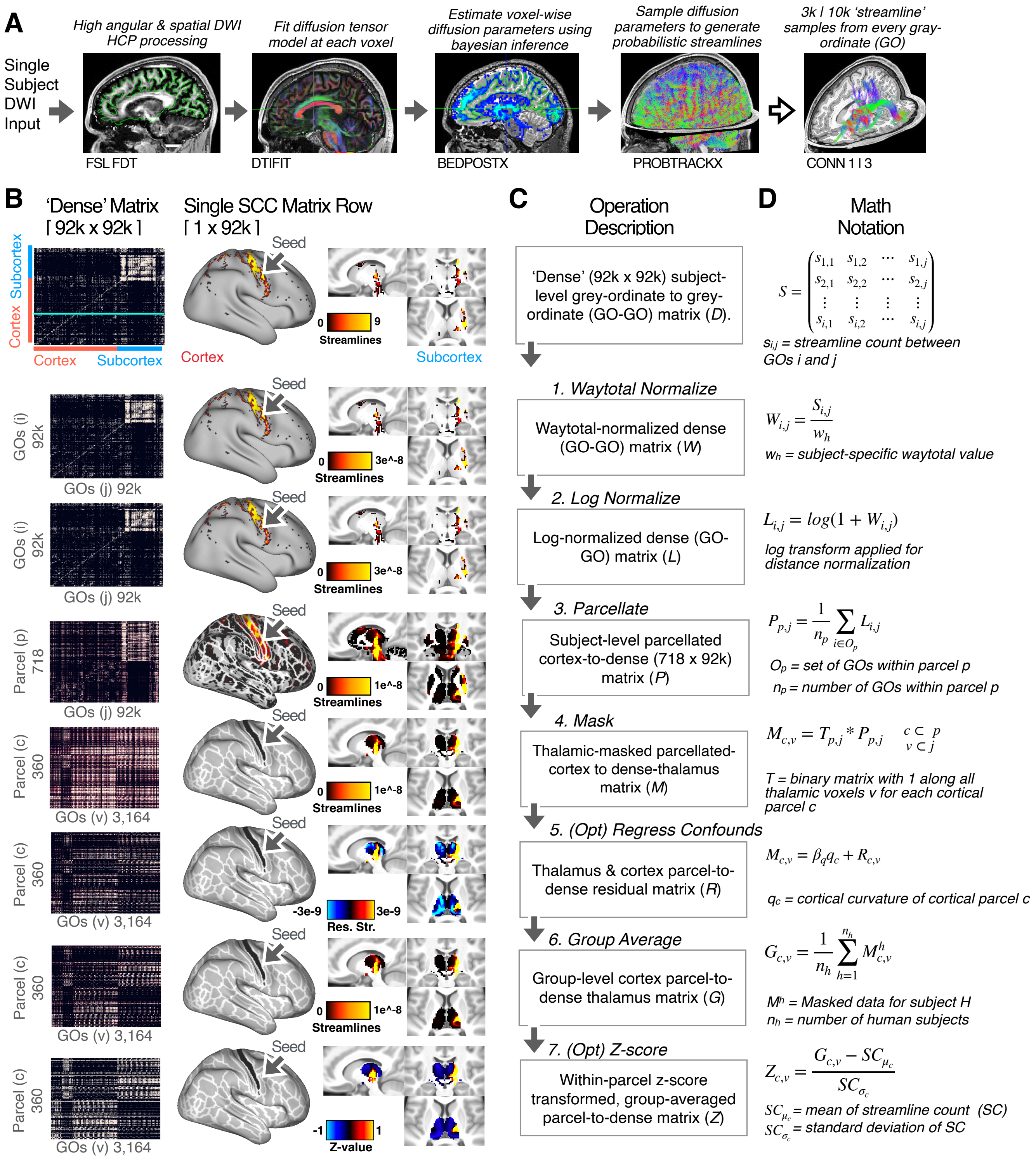


**Fig. S1.** Whole-brain dMRI-derived tractography processing pipeline. Transformation of connectivity data from raw diffusion data to a normalized, whole-brain cortical-parcel to dense-thalamus connectivity matrix. These steps include (1) waytotal normalization to account for inter-individual differences in number of streamlines generated; (2) log normalization to account for distance effects; (3) parcellation to reduce the dimensionality of the data and extract parcel by grey-ordinate streamline counts; (4) masking to extract only streamline counts in the target brain area (i.e., the thalamus); (5) optional (opt) processing step to regress out cortical curvature from streamline counts; (6) group averaging; (7) optional step to standardize the data within each cortical parcel (for visualization purposes only).

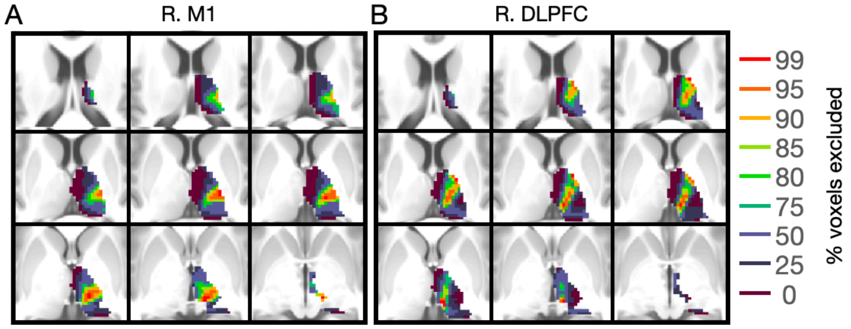


**Fig. S2.** Visualization of surviving thalamic voxels for a subset of exclusion thresholds for right **(A)** M1 and **(B)** DLPFC.

DRAFT

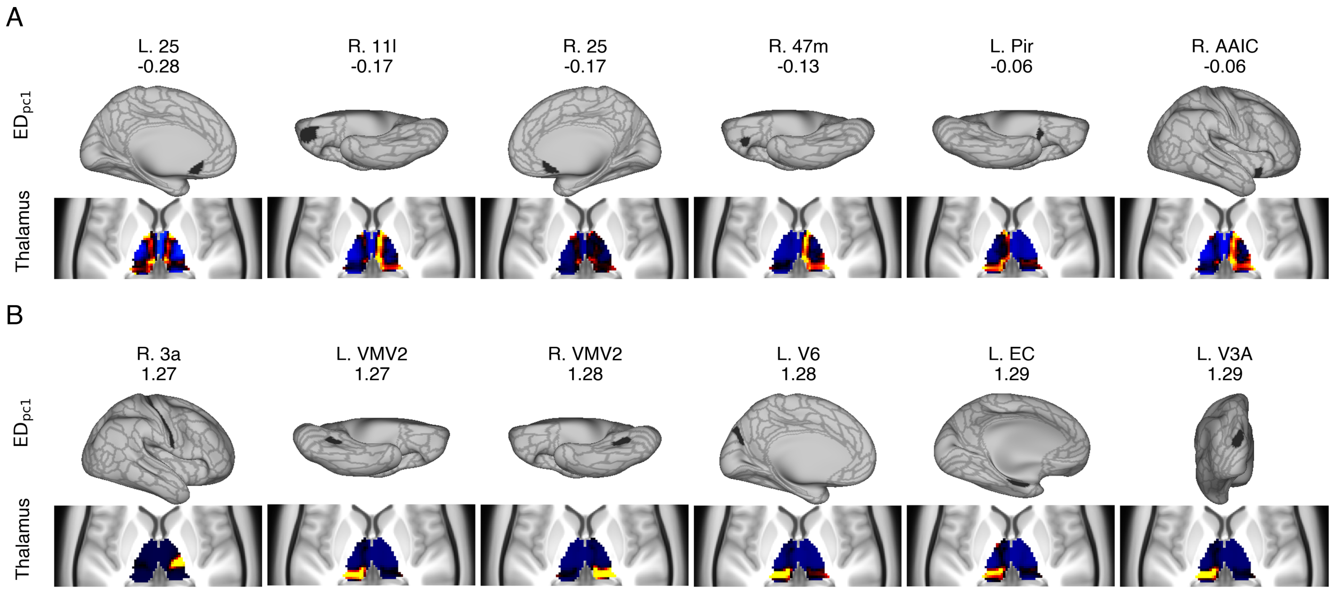


**Fig. S3.** Exemplar cortical parcels and their corresponding connectivity patterns within the thalamus. **(A)** Cortical parcels with the lowest ipsilateral $ED_{pc1}$ loadings and **(B)** highest ipsilateral $ED_{pc1}$ loadings.

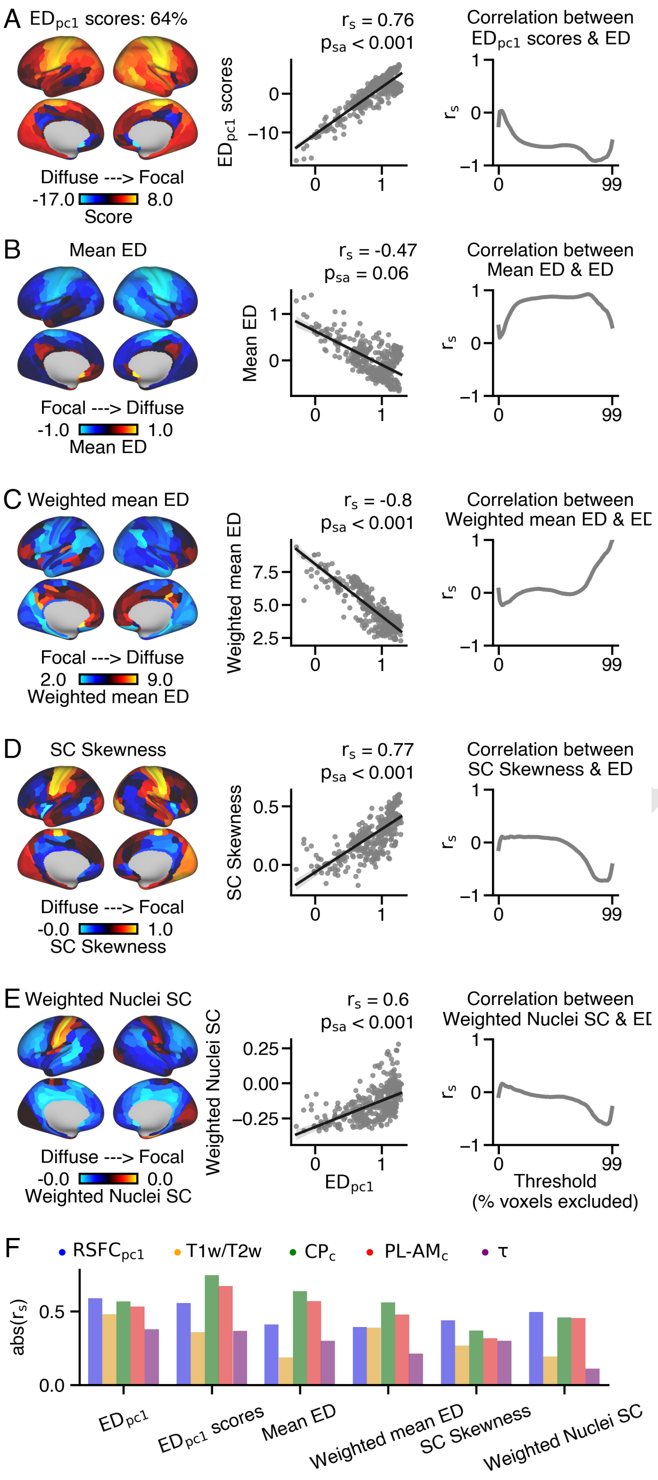


**Fig. S4.** Comparison between $ED_{pc1}$ loadings and alternative measures to quantify the extent of cortical connections within the thalamus. **(A)** $ED_{pc1}$ score cortical map (left) and comparison with $ED_{pc1}$ loadings (middle) and ED values at individual thresholds (right). PCA was applied to the transposed ED matrix (Fig. 1C) and explained 64% of the variance. **(B)** Mean ED cortical map, calculated by averaging ED across thresholds for each cortical parcel. **(C)** Weighted Mean ED cortical map, calculated by weighting thresholds and calculating the ED across thresholds. **(D)** Streamline count (SC) Skewness cortical map, calculated as the standardized difference between mean and median SC values across the thalamus. **(E)** Weighted Nuclei SC cortical map, calculated by multiplying Mean SC for each cortical parcel with the corresponding thalamic nucleus's volume. **(F)** Correlation comparison between cortical gradients and measures of the extent of connections within the thalamus. $RSFC_{pc1}$ loadings and T1w/T2w values indicate cortical hierarchy, $CP_c$ reflects anatomical overlap across the CALB1-PVALB thalamic gradient, PL-$AM_c$ values reflect anatomical overlap across the anteromedial-posterolateral thalamic gradient, and $\tau$ represents intrinsic functional timescale. Refer to the main text for further details on these cortical gradients.

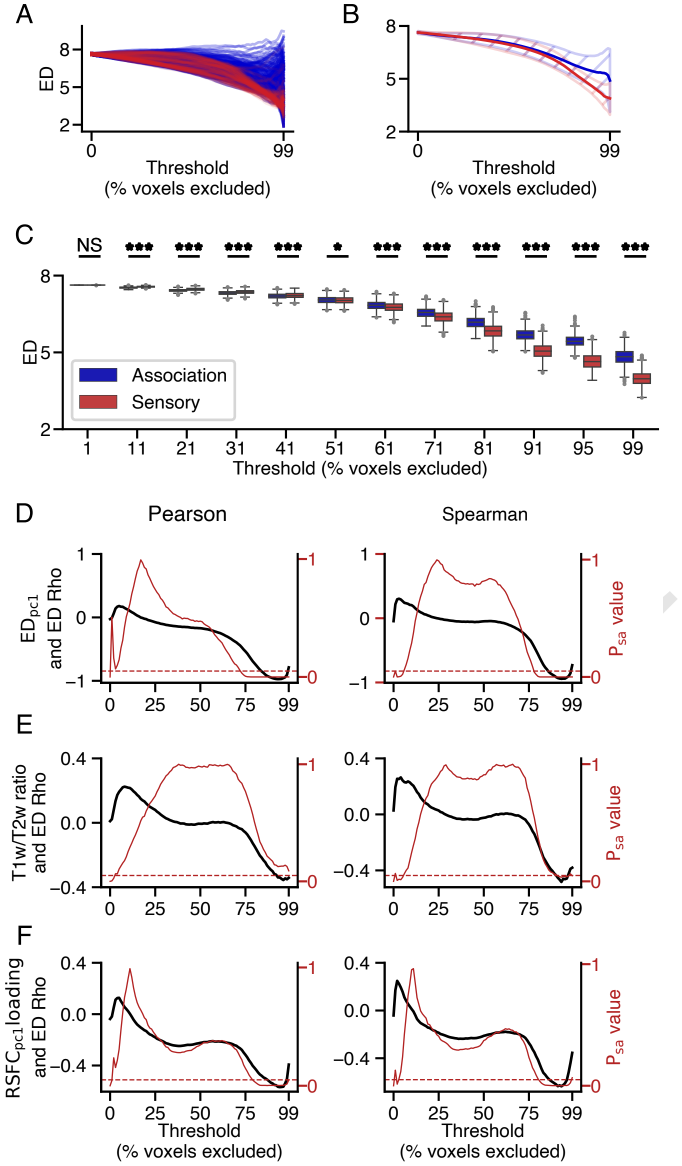


**Fig. S5.** Correspondence between cortical gradients and ED at calculated for individual thresholds. **(A-B)** Sensory (red) and association (blue) cortical parcels exhibit distinct patterns of ED values across thresholds. **(A)** shows ED values across thresholds for each cortical parcel and **(B)** shows the average ED (line) and standard deviation (shaded area) of ED across thresholds for sensory and association cortical parcels, respectively. **(C)** Sensory cortical parcels had significantly lower ED values for thresholds 99, 95, 91, 81, 71, 61, and 51, while association cortical parcels had significantly lower ED values for thresholds 41, 31, 21, and 11 (Wilcoxon rank-sum testt, $p < 0.001$). **(D)** $ED_{pc1}$ loadings significantly correlated with ED at more conservative thresholds (e.g., 73-99% (Pearson's rho) and 78-99% (Spearman's rank correlation)). The red dashed line indicates statistical significance at $p_{sa} < 0.05$. **(E)** T1w/T2w values significantly correlated with ED at more conservative thresholds (e.g., 91-99% (Spearman rank correlation)). **(F)** $RSFC_{pc1}$ loadings significantly correlated with ED at conservative thresholds (e.g., 79-99% (Pearson's correlation) and 81-98% (Spearman's rank correlation)).

$$ED_{\mu_{c,\tau_1,\tau_2}} = \frac{1}{\tau_2 - \tau_1} \sum_{t=\tau_1}^{\tau_2} ED_{c,t}$$

$$\tau_1 \leq t < \tau_2$$

$$ED_{\sigma_{c,\tau_1,\tau_2}} = \sqrt{\frac{1}{\tau_2 - \tau_1} \sum_{t=\tau_1}^{\tau_2} ED_{c,t} - ED_{\mu_{c,\tau_1,\tau_2}}}$$

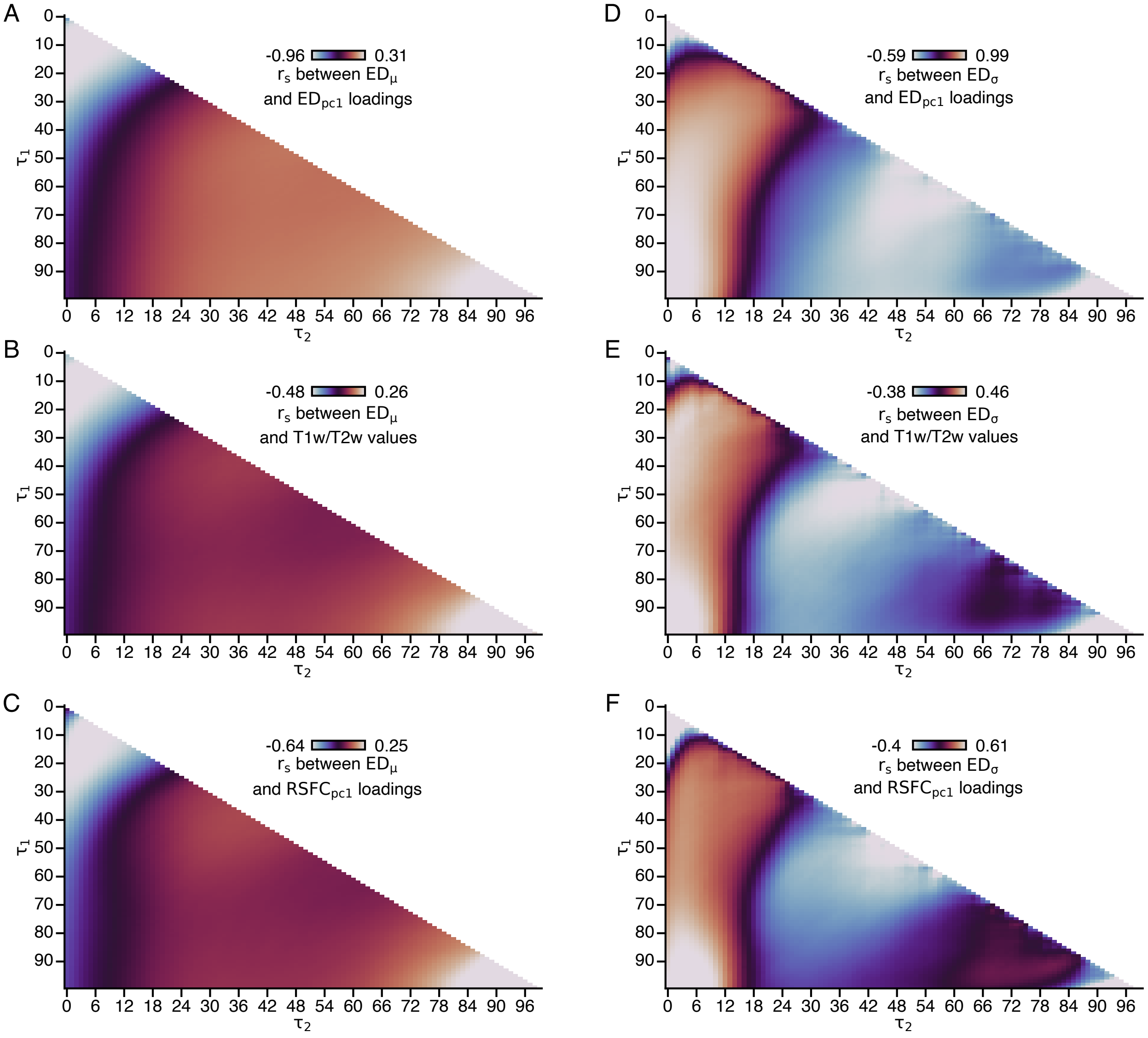


**Fig. S6.** Alternative threshold selection using range-based approaches to calculate the mean (left panels) or standard deviation (right panels) of ED within a range of thresholds. **(A-C)** Mean ED values were calculated for each cortical parcel between thresholds $\tau_1$ and $\tau_2$, and subsequently correlated with **(A)** $ED_{pc1}$ loadings, **(B)** T1w/T2w values, and **(C)** $RSFC_{pc1}$ loadings. **(D-E)** Standard deviation of ED values between thresholds $\tau_1$ and $\tau_2$ was calculated for each cortical parcel, and subsequently correlated with **(D)** $ED_{pc1}$ loadings, **(E)** T1w/T2w values, and **(F)** $RSFC_{pc1}$ loadings.

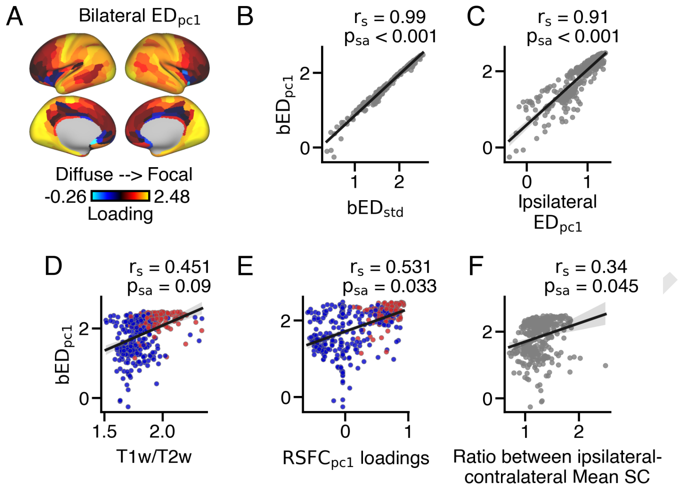


**Fig. S7.** Cortical variation in the extent of connections within bilateral thalamus. **(A)** ED was calculated for right and left thalamus for each cortical parcel for 100 thresholds. These values were summed together and the resulting bilateral ED values were used as input into PCA, yielding the first principal component, indexing the extent of anatomical connectivity across bilateral thalamus ($bED_{pc1}$). **(B)** Bilateral $ED_{pc1}$ loadings strongly correlated with Bilateral $ED_{\sigma}$ (std) values. **(C)** Bilateral $ED_{pc1}$ loadings strongly correlated with ipsilateral $ED_{pc1}$ loadings. **(D)** Bilateral $ED_{pc1}$ loadings exhibited a trending correlation with T1w/T2w values. **(E)** Bilateral $ED_{pc1}$ loadings significantly correlation with $RSFC_{pc1}$ loadings. **(F)** Bilateral $ED_{pc1}$ loadings significantly correlation with the ratio of Mean SC between ipsilateral and contralateral thalamus, reflecting the laterality of thalamic terminations. A ratio of one indicates cortical parcels with with equal terminations in left and right thalamus, while a higher positive ratio indicates cortical parcels with predominantly ipsilateral thalamic terminations.

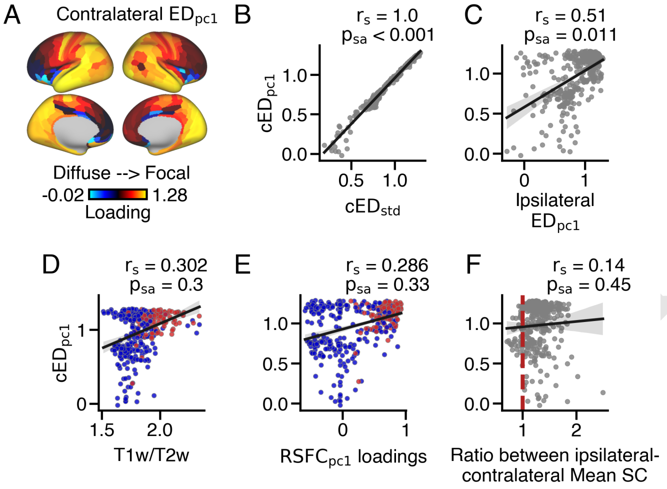


**Fig. S8.** Cortical variation in the extent of connections within contralateral thalamus. **(A)** ED was calculated to quantify the extent of each cortical parcel's connections within contralateral thalamus. These values were used as input into PCA, yielding the first principal component, indexing the extent of anatomical connectivity across contralateral thalamus ($cED_{pc1}$). **(B)** Contralateral $ED_{pc1}$ loadings strongly correlated with contralateral $cED_{\sigma}$ (std) values. **(C)** Contralateral $ED_{pc1}$ loadings significantly correlation with ipsilateral $ED_{pc1}$ loadings. **(D)** Contralateral $ED_{pc1}$ loadings did not significantly correspond with T1w/T2w values. **(E)** Contralateral $ED_{pc1}$ loadings did not significantly correspond with $RSFC_{pc1}$ loadings. **(F)** Contralateral $ED_{pc1}$ loadings did not have a significant correlation with the ratio of Mean SC between ipsilateral and contralateral thalamus, which indexes the laterality of thalamic terminations (i.e., a ratio less than one (red dashed lined) indexes cortical parcels with stronger contralateral thalamic connectivity). Some cortical parcels had stronger anatomical connections to contralateral thalamus relative to ipsilateral thalamus, and those cortical parcels had higher contralateral $ED_{pc1}$ loadings.

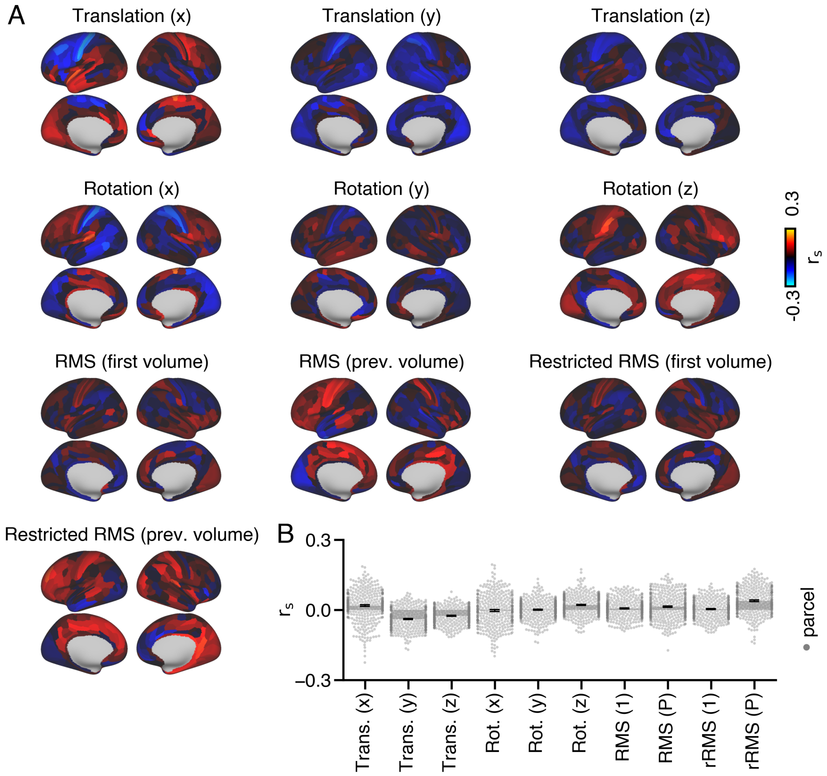


**Fig. S9.** $ED_{pc1}$ loadings do not correspond strongly with individual variation in motion. Motion parameters, including translation and rotation in x, y, and z directions, were extracted from FSL's Eddy tool. RMS (root mean square) and restricted RMS measures were obtained for 813 subjects. These estimates were averaged across volumes, resulting in 10 motion measures per subject. Correlations were then calculated between these motion measures and ipsilateral $ED_{pc1}$ loadings for each cortical parcel, separately. **(A)** This process yielded a Spearman rho value for each cortical parcel for each motion parameter. **(B)** Weak correlations were observed on average between $ED_{pc1}$ loadings and motion estimates across all cortical parcels. The mean correlation between $ED_{pc1}$ loadings and all motion parameters was close to zero.

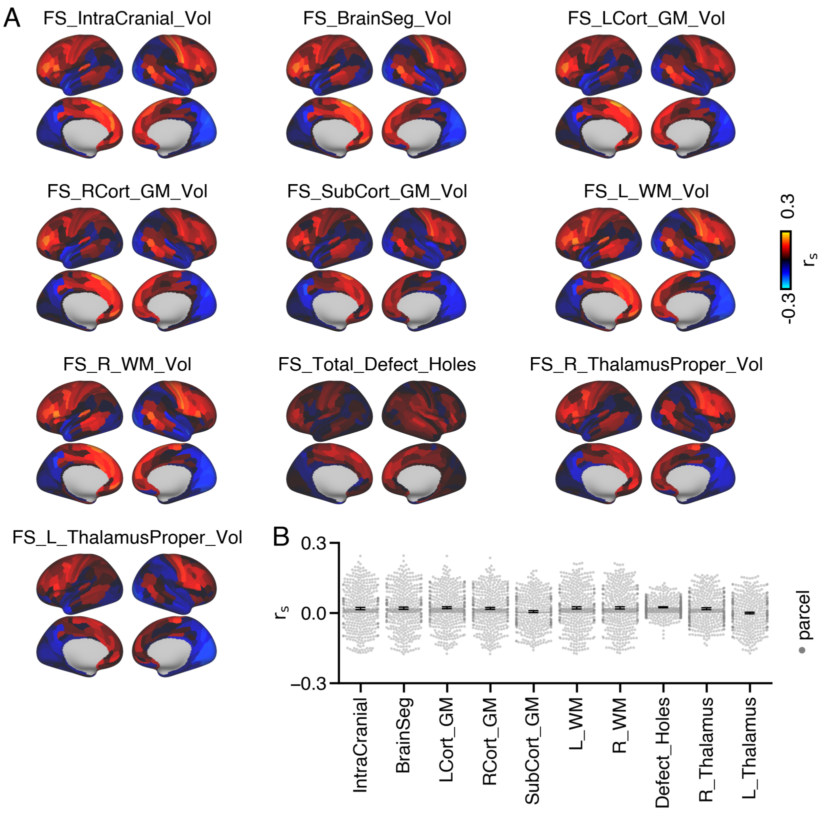


**Fig. S10.** $ED_{pc1}$ loadings do not correspond strongly with individual variation in white matter (WM), gray matter (GM), or defects. Freesurfer measures, including intracranial volume, brain segmentation volume, GM and WM volumes for left and right cortex, subcortical GM volume, total defect holes, and total volume of left and right thalamus, were obtained from the HCP's ConnectomeDB. Correlations were calculated between these 10 Freesurfer measures and ipsilateral $ED_{pc1}$ loadings for each cortical parcel, separately. **(A)** This process resulted in 360 Spearman rho values for each Freesurfer measure. **(B)** Weak correlations were observed on average between $ED_{pc1}$ loadings and Freesurfer measures across all cortical parcels. The mean correlation between $ED_{pc1}$ loadings and all Freesurfer measures was close to zero.

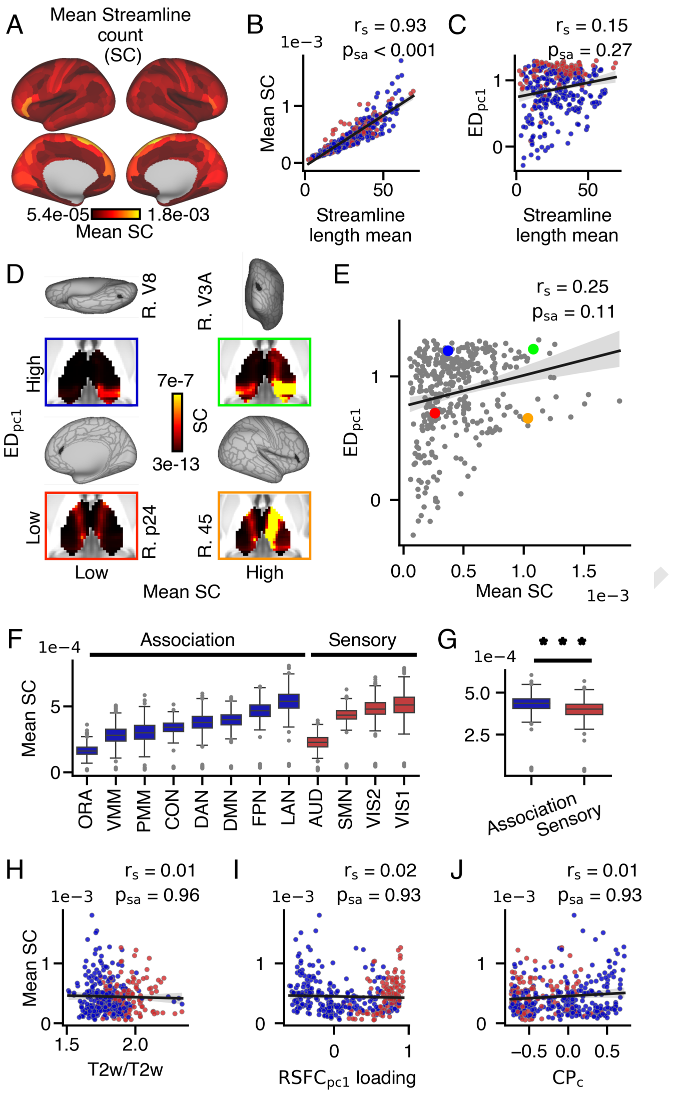


**Fig. S11.** Dissociation between extent and strength in thalamic connectivity patterns. **(A)** Mean streamline count (Mean SC) was calculated for each cortical parcel, representing the relative strength of connectivity. Parcels with higher Mean SC values have more streamline terminations within the thalamus. **(B)** Mean SC strongly correlated with mean streamline length. **(C)** Mean streamline length weakly correlated with $ED_{pc1}$. **(D-E)** Exemplar cortical parcels were selected based on their $ED_{pc1}$/Mean SC values: low/high (R. 45; orange), high/high (R. V3A; green), low/low (R. p24; red), and high/low (R. V8; blue) **(F-G)** Mean SC varied within and between cortical networks, with slightly lower values in sensory (median = 4.0e-04) compared to association (median = 4.4e04) networks (Wilcoxon signed-rank test, p = 3.9e-94). There was no correlation between Mean SC and **(H)** T1w/T2w values, **(I)** $RSFC_{pc1}$ loadings, or **(J)** $CP_c$ values.

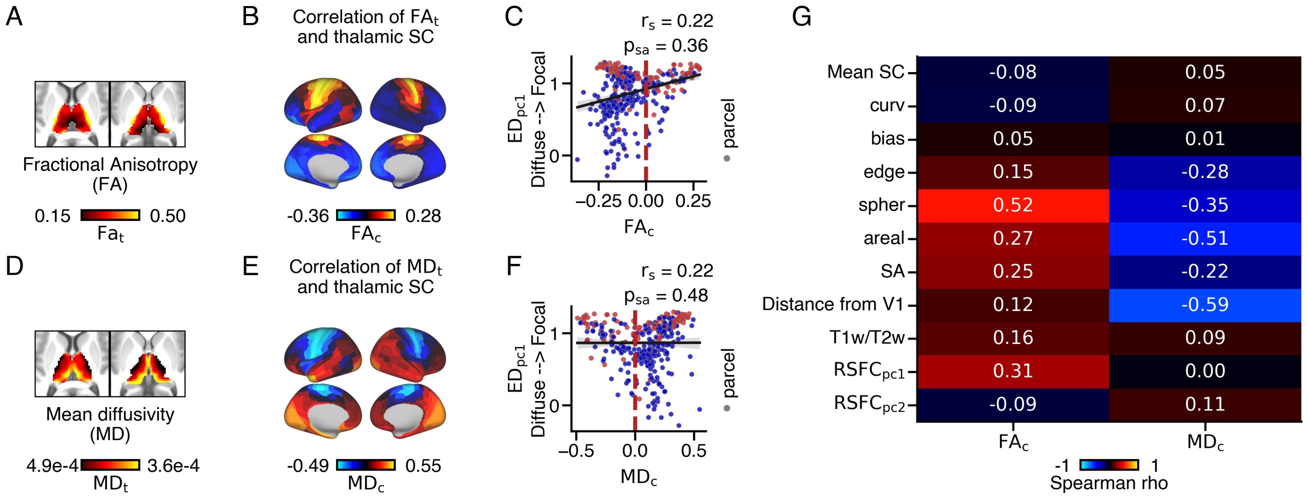


**Fig. S12.** $ED_{pc1}$ loadings do not correspond with anatomical overlap with thalamic fractional anisotropy $(FA_t)$ and mean diffusivity $(MD_t)$ at the group level. FS and MD values were obtained from FSL's DTIFIT and group-averaged across 828 HCP subjects. **(A)** Fractional anisotropy thalamic gradient $(FA_t)$. **(B)** The overlap between each cortical parcel's anatomical connectivity along the $FA_t$ gradient was calculated for each cortical parcel. Across cortex, warmer colors reflect cortical parcels that preferentially couple with thalamic areas with higher FA values $(FA_c)$. **(C)** $FA_c$ did not significantly correlate with $ED_{pc1}$ loadings. **(D)** Mean diffusivity thalamic gradient $(MD_t)$. **(E)** The overlap between each cortical parcel's anatomical connectivity along the $MD_t$ gradient was calculated for each cortical parcel. Across cortex, warmer colors reflect cortical parcels that preferentially couple with thalamic areas with higher MD values $(MD_c)$. **(F)** $MD_c$ did not significantly correlate with $ED_{pc1}$ loadings. **(G)** Correlations between $FA_t$ and $MD_t$ and other cortical gradients.

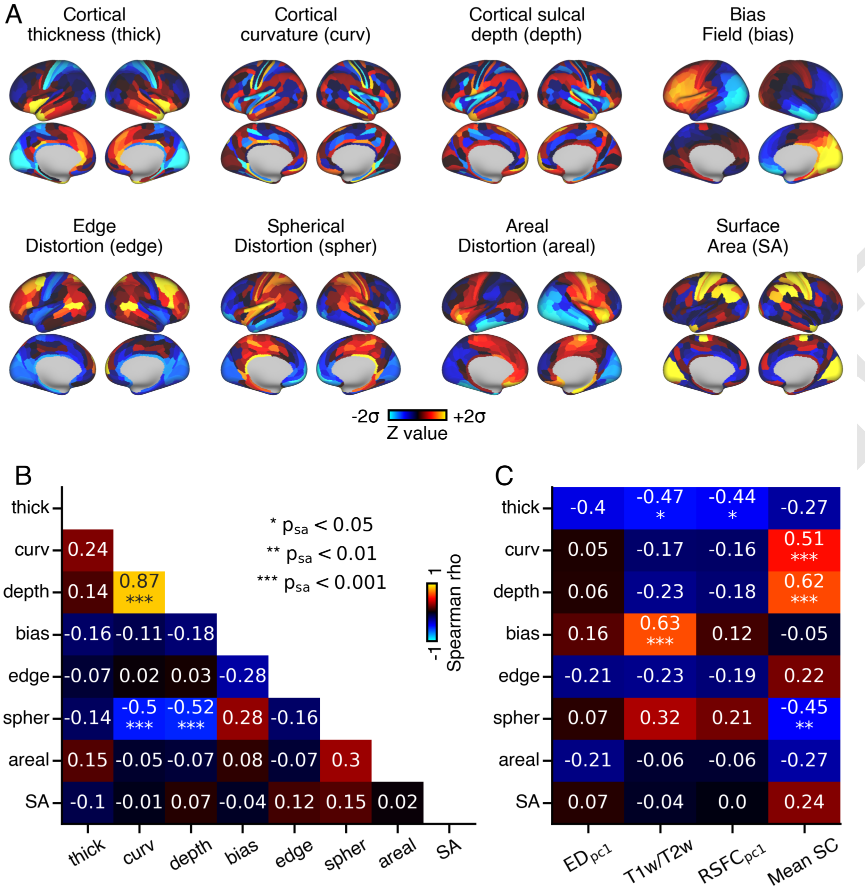


**Fig. S13.** $ED_{pc1}$ loadings do not correspond strongly with measures of cortical geometry, bias, or distortion. **(A)** Cortical maps depicting averaged cortical thickness, cortical curvature, sulcal depth, bias field, edge distortion, spherical distortion, and areal distortion values across HCP subjects. **(B)** Correlation matrix showing pairwise correlations among cortical geometry, bias, and distortion maps. **(C)** Correlations between cortical geometry, bias, and distortion maps, and $ED_{pc1}$, T1w/T2w, $RSFC_{pc1}$, and Mean SC values.

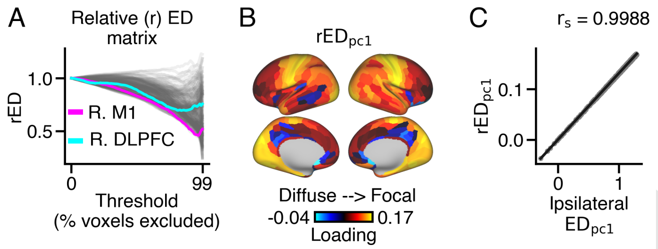


**Fig. S14.** Normalizing $ED$ values to account for size differences between left and right thalamus. **(A)** ED values were divided by the unthresholded ED value (when 0% of voxels were excluded) for each cortical parcel, creating a relative measure of ED. The unthresholded value is the same for all cortical parcels within the same hemisphere, as it reflects size differences between the left and right thalamus. **(B)** Cortical map of relative (r) $ED_{pc1}$ loadings. **(C)** $rED_{pc1}$ loadings nearly perfectly correlated with $ED_{pc1}$ loadings.

DRAFT

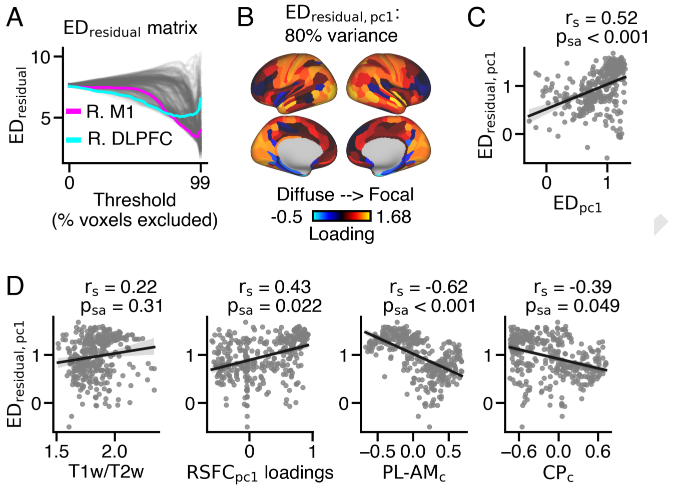


**Fig. S15.** Correspondence between $ED_{pc1}$ loadings before and after regressing out subject-level cortical curvature from subject-level Mean SC values. The residuals were then group-averaged for further analysis. **(A)** $ED_{residual}$ values were calculated between surviving thalamic voxels with the highest residual streamline count for each of 100 thresholds. **(B)** PCA was performed on the $ED_{residual}$ matrix, and the first principal component ($ED_{residual,pc1}$ loadings) accounted for 80% of the variance. **(C)** $ED_{pc1}$ loadings significantly correlated with $ED_{residual,pc1}$ loadings. **(D)** $ED_{residual,pc1}$ loadings did not significantly correlate with myelin, but did significantly correlate with $RSFC_{pc1}$ loadings, $PL\text{-}AM_c$ values, and $CP_c$ values (see main text for further details)

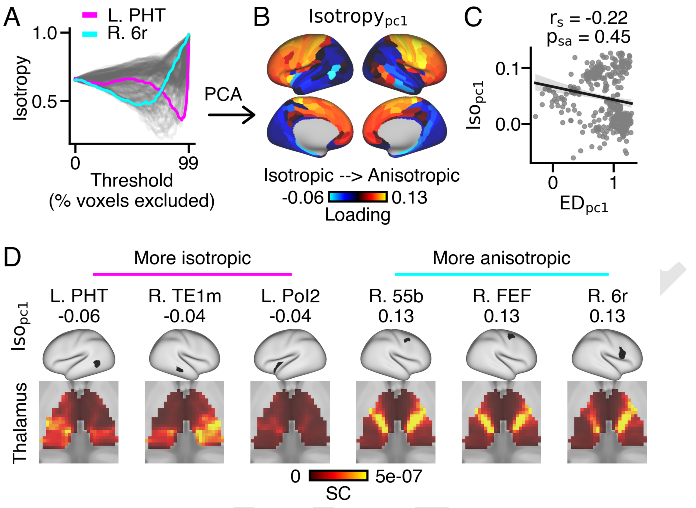


**Fig. S16.** Estimating the isotropy to examine cortical variation in the shape of thalamic connectivity patterns. **(A)** Isotropy of ipsilateral thalamic connections was calculated for each cortical parcel for 100 thresholds. Two examplar parcels (PHT, magenta; 6r, cyan) are shown. Cooler colors reflect more isotropic connectivity patterns (the connectivity pattern evenly extends in x, y, and z directions), while warmer colors reflect more anisotropic connectivity patterns (e.g., does not extend evenly in x, y, and z directions). **(B)** PCA was performed using the matrix in panel A, resulting in a single value for each cortical parcel ($Isotropy_{pc1}$ loadings). Cooler colors reflect cortical parcels with more isotropic thalamic connectivity patterns, while warmer colors reflect cortical parcels with more anisotropic thalamic connectivity patterns. **(C)** $ED_{pc1}$ loadings did not strongly correspond with $Isotropy_{pc1}$ loadings. **(D)** Exemplar thalamic connectivity patterns for six cortical parcels with highly isotropic (left 3 panels) or highly anisotropic (right 3 panels) thalamic connectivity patterns.

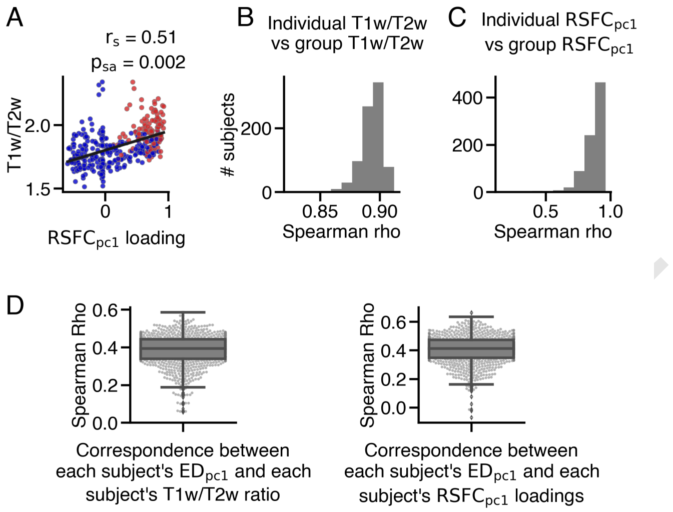


**Fig. S17.** Comparison of group-averaged and individual-level T1w/T2w values and $RSFC_{pc1}$ loadings. **(A)** Group-averaged T1w/T2w values significantly correlated with $RSFC_{pc1}$ loadings. Sensory cortical parcels are represented by red dots, while association cortical parcels are represented by blue dots. **(B)** Individual subjects' T1w/T2w values highly correlated with group-averaged T1w/T2w values (mean: 0.89, median: 0.89, SEM = 0.0003, STD = 0.009). **(C)** Individual subjects' $RSFC_{pc1}$ loadings highly correlated with group-averaged $RSFC_{pc1}$ loadings (mean: 0.87, median: 0.89, SEM = 0.0003, STD = 0.08). **(D)** Each subject's $ED_{pc1}$ loadings were correlated with their individual T1w/T2w and $RSFC_{pc1}$ values. On average, individual subjects' $ED_{pc1}$ loadings modestly correlated with their respective T1w/T2w values across cortex (mean: 0.38, median: 0.39, SEM = 0.003, STD = 0.08) and $RSFC_{pc1}$ loadings (mean: 0.40, median: 0.41, SEM = 0.003, STD = 0.10).

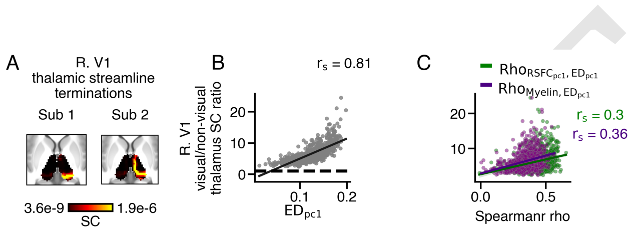


**Fig. S18.** Inter-subject variability of the extent of cortical connections within the thalamus. **(A)** Exemplar thalamic connectivity patterns from right V1 for two subjects, showing focal (sub 1) and diffuse (sub 2) patterns. While both subjects showed terminations in the visual thalamus (e.g., pulvinar and lateral geniculate nucleus), subject 2 displayed additional terminations extending to the anterior thalamus, potentially indicating false positives (131). **(B)** The ratio of streamline counts from right V1 to visual thalamus (pulvinar and lateral geniculate nucleus) relative to non-visual thalamus (all other thalamic nuclei) highly correlated with right V1 $ED_{pc1}$ loadings across subjects. **(C)** The ratio of right V1 streamline terminations to visual and non-visual thalamus modestly corresponded with rho values between $ED_{pc1}$ and T1w/T2w (purple) and $RSFC_{pc1}$ (green) values.

DRAFT

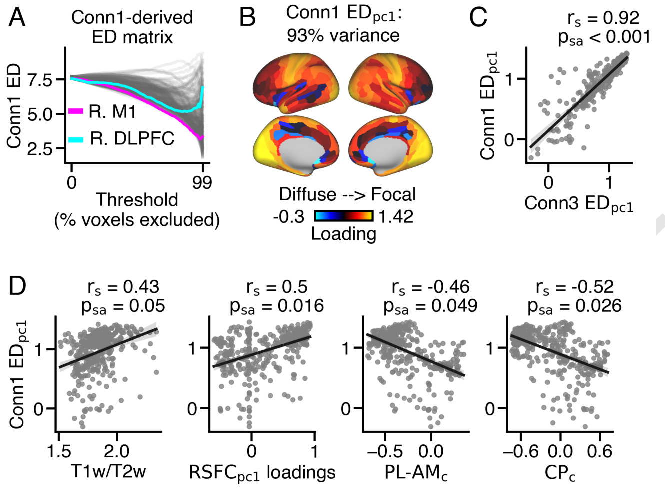


**Fig. S19.** Comparison of $ED_{pc1}$ loadings derived from bidirectional white-ordinate to gray-ordinate (Conn3) and unidirectional gray-ordinate to gray-ordinate (Conn1) tractography seeding strategies. **(A)** ED matrix derived from Conn1 tractography seeding strategy. **(B)** $ED_{pc1}$ loading map derived from Conn1 seeding strategy. PCA was performed on the Conn1-derived ED matrix, resulting in 360 loadings from the first principal component (PC1), which accounted for 93% of the variance. **(C)** Conn1 and Conn3-derived $ED_{pc1}$ loadings strongly corresponded with one another. **(D)** Conn1-derived $ED_{pc1}$ loadings significantly correlated with T1w/T2w, $RSFC_{pc1}$, $PL\text{-}AM_c$, and $CP_c$ values.

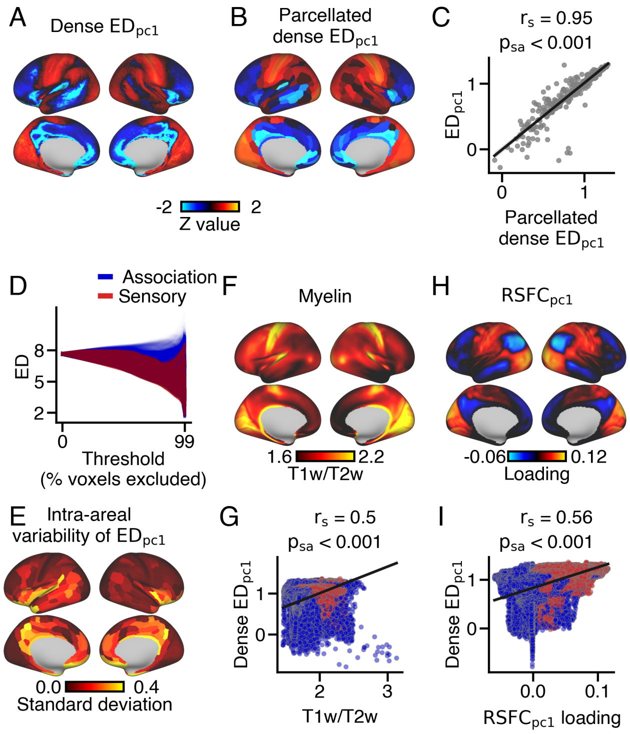


**Fig. S20.** $ED_{pc1}$ loadings on dense (i.e., grey-ordinate) cortical data replicate observations from parcellated cortical data. **(A)** Cortical map of $ED_{pc1}$ loadings for each of the 59,412 cortical vertices (Dense $ED_{pc1}$). **(B)** Dense $ED_{pc1}$ loadings were averaged within 360 cortical parcels (i.e., parcellated) from Glasser et al. (45) (Parcellated Dense $ED_{pc1}$). **(C)** Parcellated Dense $ED_{pc1}$ loadings strongly correlated with $ED_{pc1}$ loadings calculated from parcel-level data. **(D)** ED matrix derived from group-averaged dense cortical data, showing ED values for each cortical vertex across 100 thresholds. **(E)** Variation of dense $ED_{pc1}$ loadings within each cortical parcel. **(F)** Dense cortical myelin map, calculated by averaging T1w/T2w values across 828 subjects. Sensory cortical vertices have higher T1w/T2w values compared to association cortical vertices. **(G)** Dense $ED_{pc1}$ loadings significantly correlated with dense T1w/T2w values across the cortex. **(H)** Loading map of the dense cortical principal functional resting-state gradient ($RSFC_{pc1}$). Sensory cortical vertices have higher loadings compared to association cortical vertices. **(I)** $ED_{pc1}$ loadings significantly correlated with $RSFC_{pc1}$ loadings across the cortex.

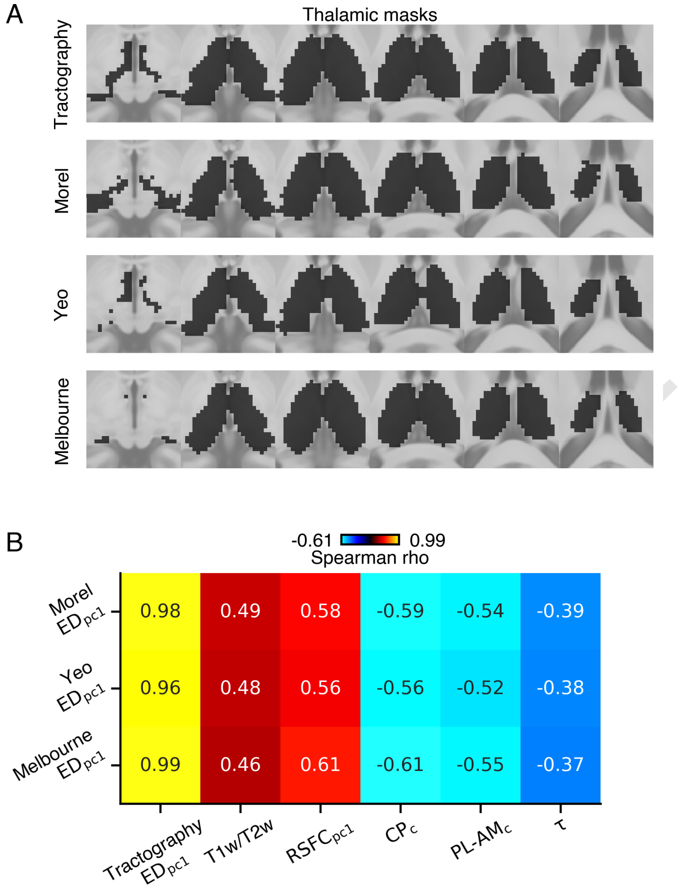


**Fig. S21.** Comparison of ipsilateral $ED_{pc1}$ loadings calculated using different thalamic masks. **(A)** Binary thalamic masks: the top panel shows the mask used for tractography seeding, while the subsequent panels show thalamic masks derived from the Morel atlas (57), the Yeo atlas (119), and the Melbourne atlas (120). **(B)** $ED_{pc1}$ loadings derived from each thalamic mask exhibited high similarity to the tractography mask, as well as consistent relationships with the T1w/T2w, $RSFC_{pc1}$, $CP_c$, $PL\text{-}AM_t$, and $\tau$ cortical gradients.

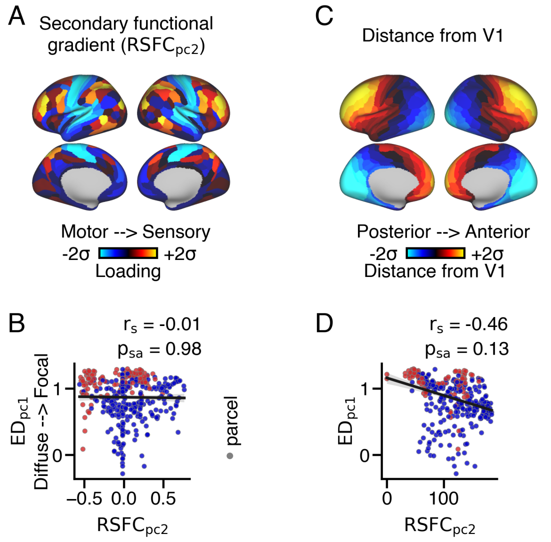


**Fig. S22.** Inter-cortical variation of the extent of connections within the thalamus is specific to the sensory-association hierarchical cortical gradient. **(A)** Cortical map showing the loading values for the secondary functional resting-state gradient ($RSFC_{pc2}$), obtained through PCA of group-averaged resting-state functional connectivity data. Sensory cortical parcels have higher loadings compared to association and motor cortical parcels. **(B)** No significant correlation was observed between $ED_{pc1}$ loadings and $RSFC_{pc2}$ loadings. **(C)** Anteroposterior cortical gradient quantified by distance from visual area 1 (V1). Anterior cortical parcels had higher values compared to posterior cortical parcels. **(D)** $ED_{pc1}$ loadings did not significantly correlate with distance from V1.

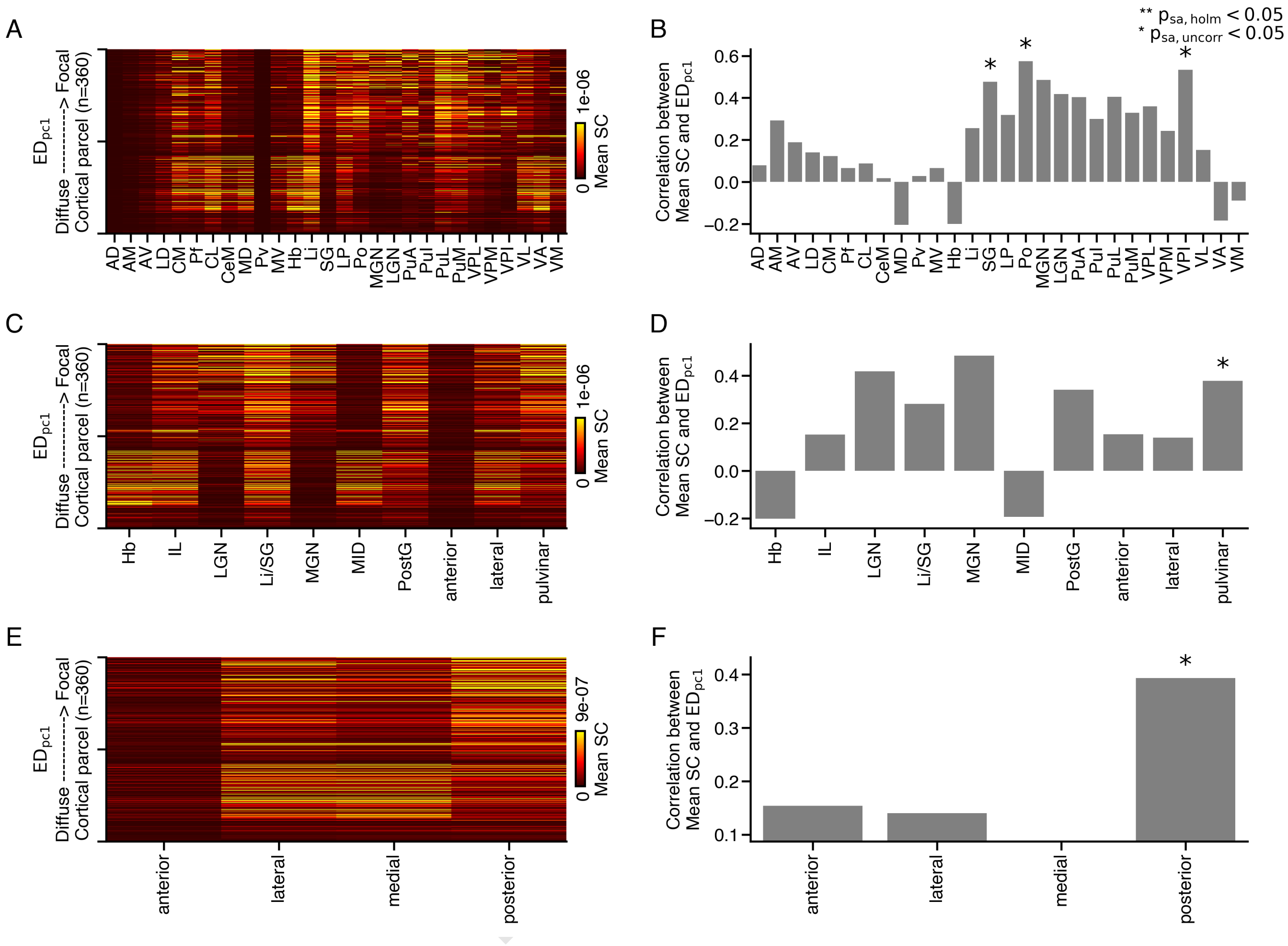


**Fig. S23.** **(A)** Streamline count (SC) connectivity matrix between each cortical parcel and each thalamic nucleus. **(B)** Correlations between $ED_{pc1}$ loadings and Mean SC for each thalamic nucleus. Cortical parcels with focal thalamic connections exhibited preferential coupling to SG, Po, and VPI nuclei, but these relationships did not survive correction for multiple comparisons. **(C)** SC connectivity matrix between each cortical parcel and minor thalamic subdivisions. **(D)** Correlations between $ED_{pc1}$ loadings and Mean SC within each minor subdivision of the thalamus. Cortical parcels with focal thalamic connections exhibited preferential coupling to the pulvinar, but this relationship did not survive correction for multiple comparisons. **(E)** SC connectivity matrix between each cortical parcel and major thalamic subdivisions. **(F)** Correlation between $ED_{pc1}$ loadings and Mean SC within each major subdivision of the thalamus. Cortical parcels with focal thalamic connections exhibited preferential coupling to posterior nuclei, but this relationship did not survive correction for multiple comparisons.

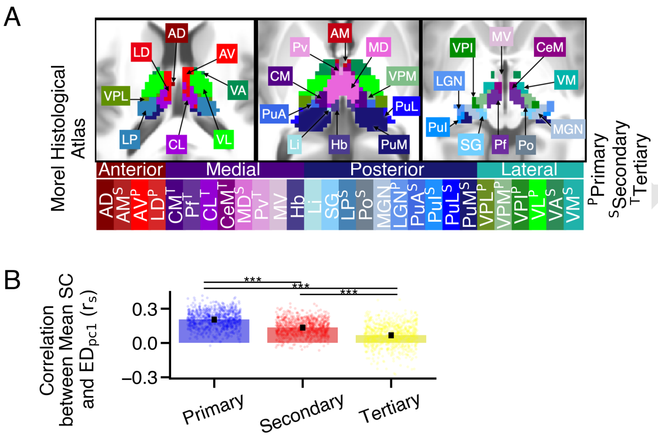


**Fig. S24.** Correspondence between Mean SC and $ED_{pc1}$ for thalamic subclasses defined in Phillips et al. (2019). Thalamic nuclei were categorized into primary, secondary, tertiary, anterodorsal, and reunions subgroups based on gene expression data (59). **(A)** Morel thalamic nuclei were assigned to primary, secondary, or tertiary subgroups. **(B)** There was a significant difference in the correlations between $ED_{pc1}$ loadings and Mean SC for primary (median: 0.21), secondary (median: 0.14), and tertiary (median: 0.07) thalamic nuclei ($\chi^2$ (3) = 1165, p < .001). Post-hoc Nemenyi tests showed significant differences for all group comparisons (*** p = 0.001).

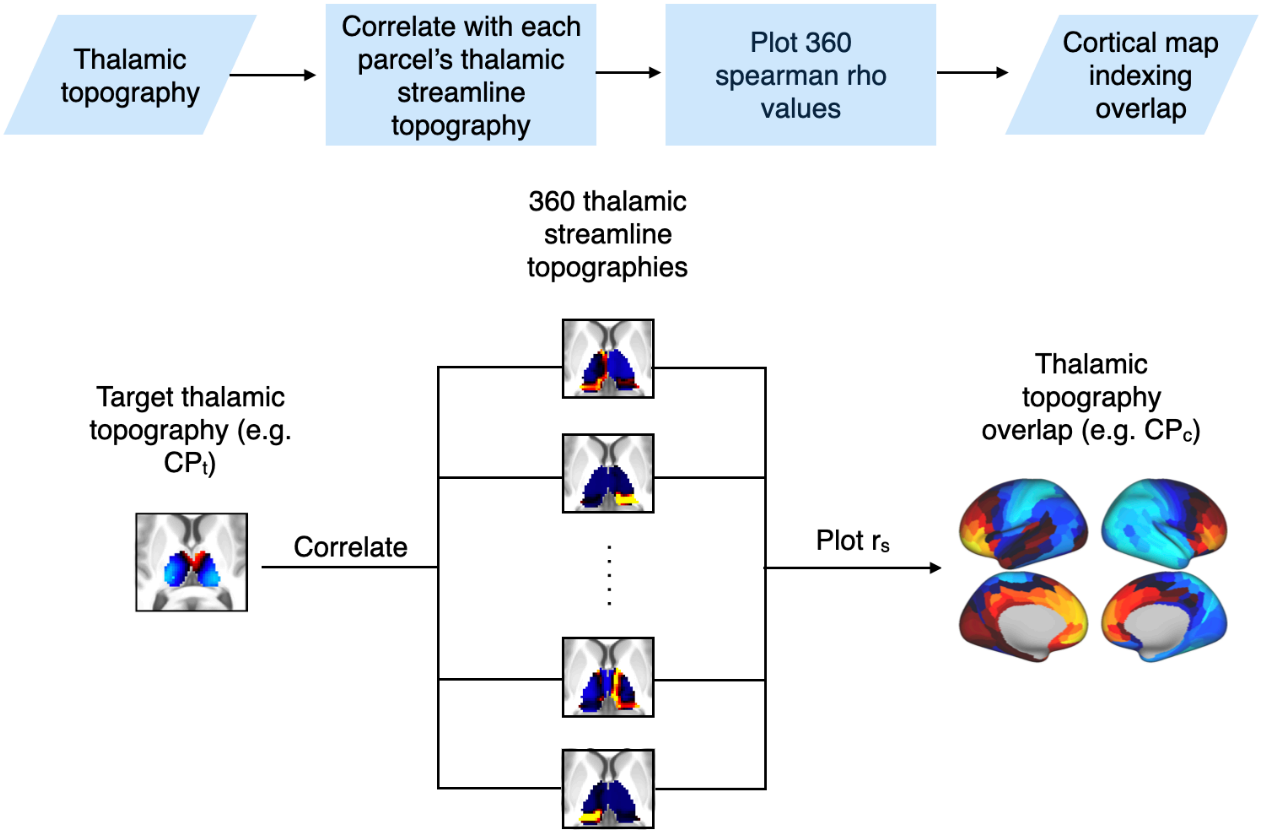


**Fig. S25.** Flow diagram (top panel) and workflow visualization (bottom panel) showing the workflow to quantify the overlap between thalamic anatomical connectivity and thalamic gradients. The streamline counts of each cortical parcel within the thalamus were correlated with a target thalamic map, yielding a cortical map of Spearman rho values. Higher rho values indicate a stronger correspondence between a cortical parcel's thalamic anatomical connectivity patterns and positive values in the target thalamic map.

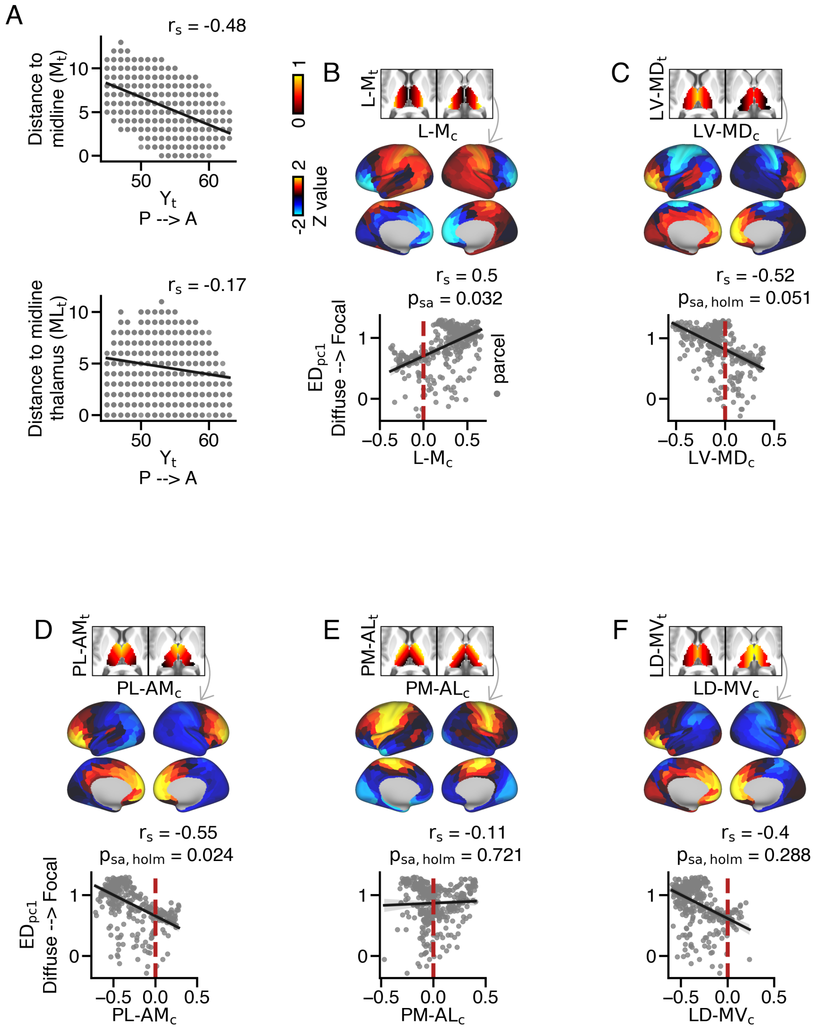


**Fig. S26.** Confounding effect of anteroposterior axis on distance to midline when indexing the mediolateral thalamic gradient. **(A)** Position along the anteroposterior $(y_t)$ thalamic axis showed stronger correspondence with distance from midline compared to distance from thalamic midline. **(B)** There was a significant correlation between the overlap of cortical parcel's thalamic anatomical connectivity and distance to midline with $ED_{pc1}$ loadings. **(C-F)** The overlap of cortical parcel's anatomical connectivity with thalamic spatial gradients using distance to midline as an index showed strong correspondence with **(C)** mediodorsal and **(D)** anteromedial thalamic gradients. However, **(E)** anterolateral and **(F)** medioventral thalamic gradients did not significantly correspond with $ED_{pc1}$ loadings.

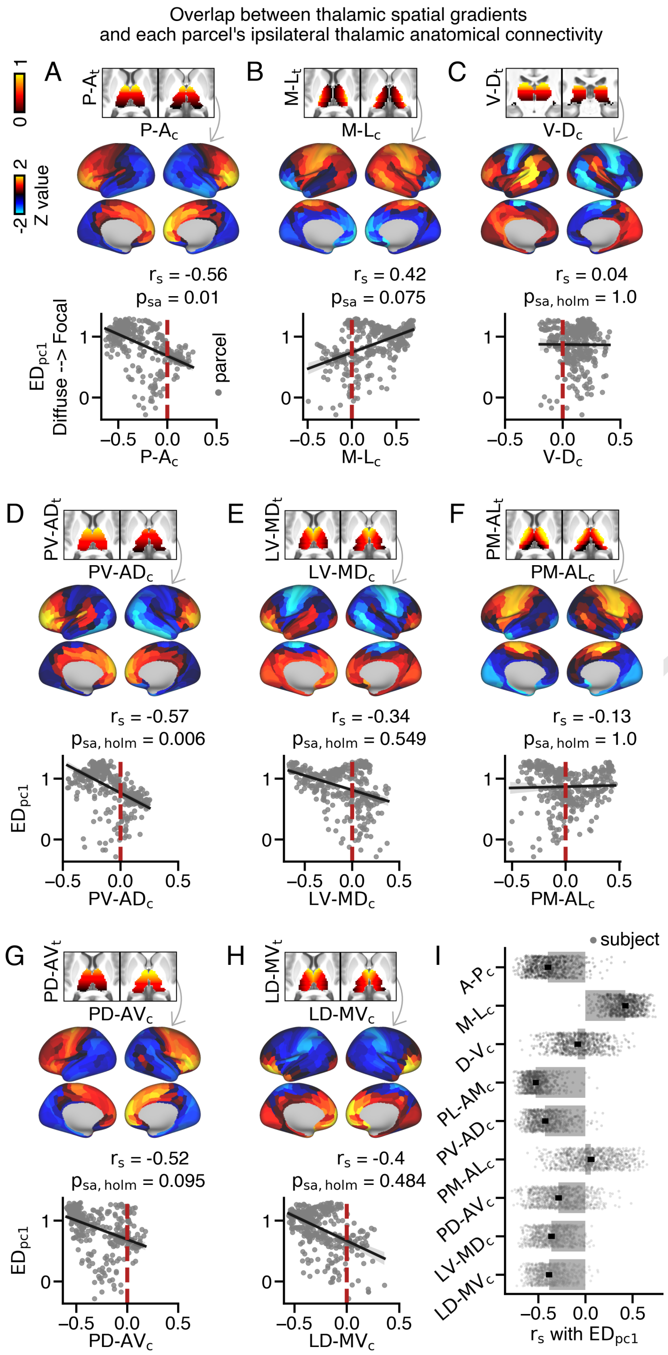


**Fig. S27.** Correspondence between ipsilateral $ED_{pc1}$ loadings and thalamic spatial gradients. **(A-H)** Overlap between each cortical parcel's thalamic anatomical connectivity patterns and the following thalamic gradients: **(A)** Anteroposterior ($P$-$A$), **(B)** Mediolateral ($M$-$L$), **(C)** Dorsoventral ($D$-$V$), **(D)** Posteroventral-Anterodorsal ($PV$-$AD$), **(E)** Lateroventral-Mediodorsal ($LV$-$MD$), **(F)** Posteromedial-Anterolateral ($PM$-$AL$), **(G)** Posterodorsal-Anteroventral ($PD$-$AV$), and **(H)** Laterodorsal-Medioventral ($LD$-$MV$). **(I)** Correlations between $ED_{pc1}$ loadings and thalamic spatial gradients across individual subjects: $A$-$P_c$ (median: -0.42), $M$-$L_c$ (median: 0.43), $D$-$V_c$ (median: -0.09), $PL$-$AM_c$ (median: -0.54), $PV$-$AD_c$ (median: -0.45), $PM$-$AL_c$ (median: 0.05), $PD$-$AV_c$ (median: -0.31), $LV$-$MD_c$ (median: -0.37), $LD$-$MV_c$ (median: -0.40).

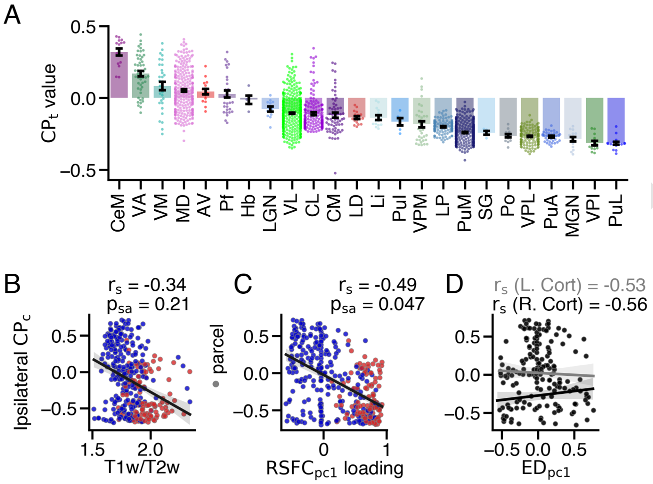


**Fig. S28.** Correspondence between cortical $CP_c$ values and cortical gradients. **(A)** Distribution of $CP_t$ values for each thalamic nucleus. **(B)** $CP_c$ values across cortex did not correspond with T1w/T2w values. **(C)** $CP_c$ values across cortex strongly correlated with $RSFC_{pc1}$ loadings. **(D)** $CP_c$ values strongly corresponded with $ED_{pc1}$ loadings in both the left and right cortex.

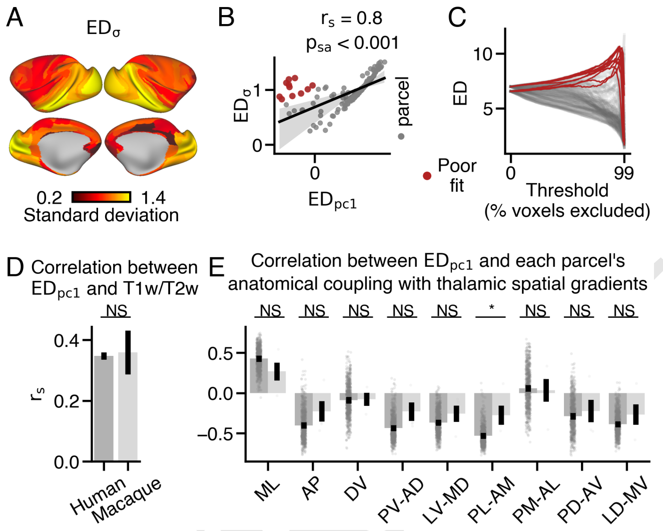


**Fig. S29.** Comparisons between human and macaque cortical connectivity patterns. **(A)** Macaque cortical map showing the standard deviation of ED ($ED_{\sigma}$) across thresholds for each cortical parcel. **(B)** $ED_{pc1}$ loadings strongly correlated with $ED_{\sigma}$ values. Some cortical parcels (red dots) where poorly fit by the linear model. **(C)** ED matrix highlighting cortical parcels with poor correspondence between $ED_{pc1}$ and $ED_{\sigma}$ values. **(D)** Comparison of the correlation between ipsilateral $ED_{pc1}$ loadings and T1w/T2w values between humans and macaques, showing no significant difference between species. **(E)** Comparison of the correlation between ipsilateral $ED_{pc1}$ loadings and each thalamic spatial gradient between humans and macaques. There was a significant main effect of gradient and species, but no interaction. A significant difference was observed in the correlation with the PL-AM thalamic gradient between humans and macaques (* $p < .05$; NS not significant). No other significant differences were found between species.

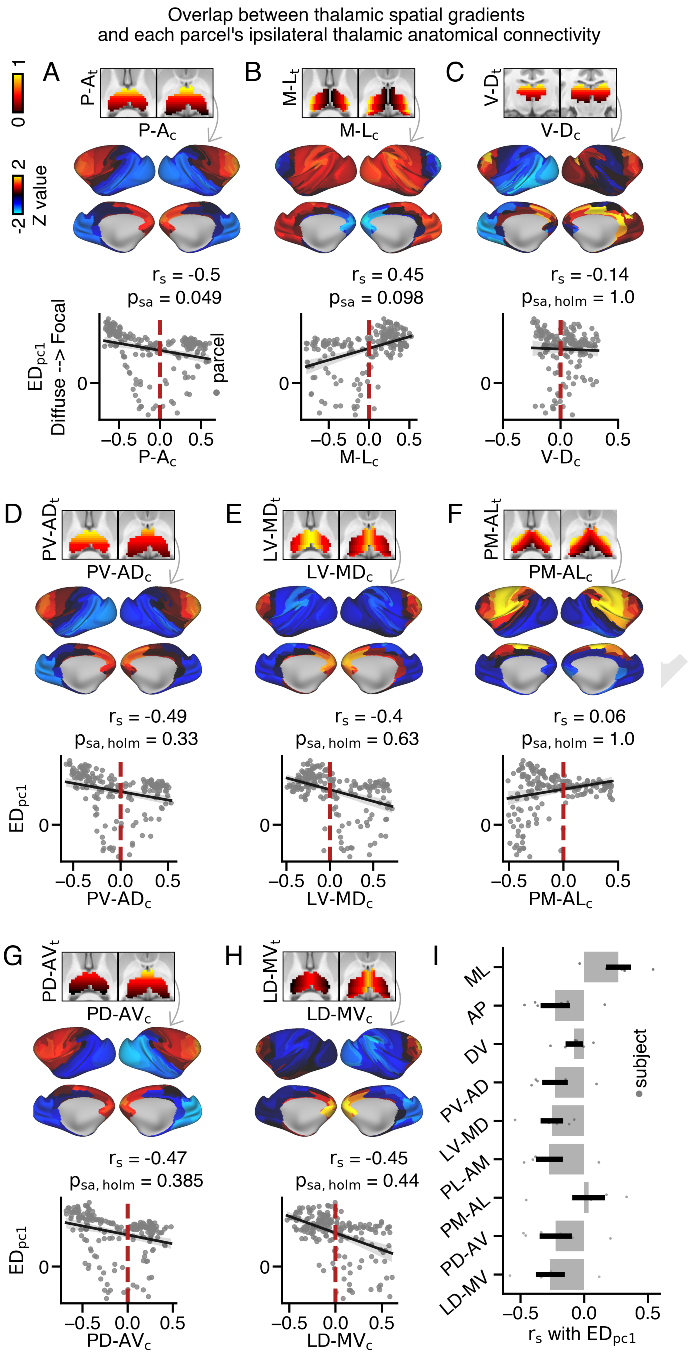


**Fig. S30.** Overlap between each cortical parcel's thalamic anatomical connectivity and the following macaque spatial thalamic gradients: **(A)** Anteroposterior ($P$-$A$) **(B)** Mediolateral ($M$-$L$) **(C)** Dorsoventral ($D$-$V$) **(D)** Posteroventral-Anterodorsal ($PV$-$AD$) **(E)** Lateroventral-Mediodorsal ($LV$-$MD$) **(F)** Posteromedial-Anterolateral ($PM$-$AL$) **(G)** Posterodorsal-Anteroventral ($PD$-$AV$) **(H)** Laterodorsal-Medioventral ($LD$-$MV$) thalamic gradients. **(I)** Median correlation between $ED_{pc1}$ loadings and thalamic spatial gradients across individual macaque subjects: $P$-$A_c$ (median: -0.27), $M$-$L_c$ (median: 0.31), $D$-$V_c$ (median: -0.06), $PL$-$AM_c$ (median: -0.33), $PV$-$AD_c$ (median: -0.27), $PM$-$AL_c$ (median: 0.08), $PD$-$AV_c$ (median: -0.26), $LV$-$MD_c$ (median: -0.24), $LD$-$MV_c$ (median: -0.29).

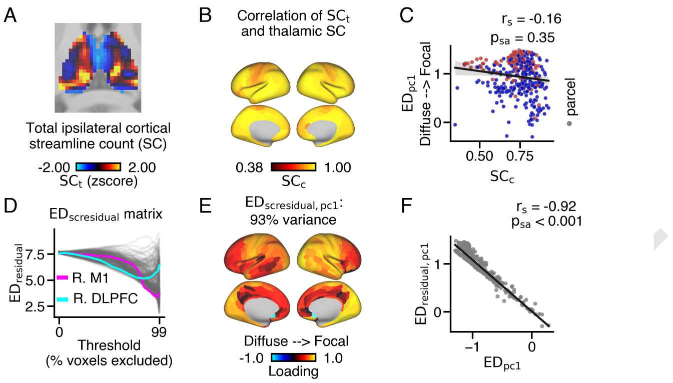


**Fig. S31.** Correspondence between group-level $ED_{pc1}$ loadings before and after regressing out each thalamic voxel's total streamline count to all ipsilateral cortical regions ($SC_t$). **(A)** For each thalamic voxel, the sum of its streamline counts (SC)s to each cortical area was calculated, showing some areas of the thalamus have stronger total structural connectivity to ipsilateral cortex. **(B)** Cortical map reflecting each cortical area's anatomical overlap across $SC_t$ values ($SC_c$). Warmer colors reflect preferential coupling to areas of the thalamus with strong connectivity to the ipsilateral cortex and cooler colors reflect preferential coupling to areas of the thalamus with weak overall connectivity to ipsilateral cortex. **(C)** $ED_{pc1}$ loadings and $SC_c$ values are only weakly correlated with one another. **(D)** After regression $SC_t$ values from each cortical area's streamline counts across ipsilateral thalamus, $ED_{scresidual}$ values were calculated between surviving thalamic voxels with the highest residual streamline count for each of 100 thresholds. **(E)** PCA was performed on the $ED_{scresidual}$ matrix, and the first principal component ($ED_{scresidual,pc1}$ loadings) accounted for 93% of the variance. **(F)** $ED_{pc1}$ loadings significantly correlated with $ED_{residual,pc1}$ loadings.

**Table 1.** Morel thalamic nuclei names, abbreviations, classes, and volumes

| Group | Name | Abbr. | Higher/First order | Phillips et al. (59) subgroup | Volume (# 2mm voxels) |
|---|---|---|---|---|---|
| Anterior group | | | | | |
| | Anterior dorsal | AD | FO | AD | 1 |
| | Anterior medial | AM | HO | Secondary | 15 |
| | Anterior ventral | AV | FO | Primary | 78 |
| | Lateral dorsal | LD | FO | Primary | 44 |
| Medial group | | | | | |
| | Centre median | CM | HO | Tertiary | 64 |
| | Parafascicular | Pf | HO | Tertiary | 73 |
| | Central lateral | CL | HO | Tertiary | 189 |
| | Central medial | CeM | HO | Tertiary | 56 |
| | Mediodorsal | MD | HO | Secondary | 310 |
| | Paraventricular | PV | HO | Tertiary | 2 |
| | Medioventral | MV | HO | Re | 9 |
| | Habenular | Hb | | | 22 |
| Posterior group | | | | | |
| | Limitans | Li | | | 27 |
| | Suprageniculate | SG | | | 18 |
| | Lateral posterior | LP | HO | Secondary | 138 |
| | Posterior | Po | HO | Secondary | 13 |
| | Medial geniculate | MGN | FO | | 70 |
| | Lateral geniculate | LGN | FO | Primary | 87 |
| | Anterior pulvinar | PuA | HO | Secondary | 34 |
| | Inferior pulvinar | PuI | HO | Secondary | 18 |
| | Lateral pulvinar | PuL | HO | Secondary | 57 |
| | Medial pulvinar | PuM | HO | Secondary | 530 |
| Lateral group | | | | | |
| | Ventral posterior lateral | VPL | FO | Primary | 142 |
| | Ventral posterior medial | VPM | FO | Primary | 55 |
| | Ventral posterior inferior | VPI | FO | Primary | 22 |
| | Ventral lateral | VL | FO | Primary | 459 |
| | Ventral medial | VA | HO | Secondary | 192 |
| | Ventral anterior | VM | HO | Secondary | 51 |

DRAFT
